# Rtt105 configurationally staples RPA and blocks facilitated exchange and interactions with RPA-interacting proteins

**DOI:** 10.1101/2022.02.05.479199

**Authors:** Sahiti Kuppa, Jaigeeth Deveryshetty, Rahul Chadda, Jenna Mattice, Nilisha Pokhrel, Vikas Kaushik, Angela Patterson, Nalini Dhingra, Sushil Pangeni, Marisa K. Sadauskas, Sajad Shiekh, Hamza Balci, Taekjip Ha, Xiaolan Zhao, Brian Bothner, Edwin Antony

**Affiliations:** Department of Biochemistry, Saint Louis University School of Medicine, St. Louis, MO 63104; Department of Chemistry and Biochemistry, Montana State University, Bozeman, MT 59717; Department of Biological Sciences, Marquette University, Milwaukee, WI 53201; Molecular Biology Department, Memorial Sloan Kettering Cancer Center, New York, NY 10065; Department of Biophysics and Biophysical Chemistry, Johns Hopkins University School of Medicine, Baltimore, MD 21205; Department of Physics, Kent State University, Kent, OH 44242

**Keywords:** RPA, DNA Repair and Recombination, HDX-MS, DNA binding, Rtt105

## Abstract

Replication Protein A (RPA) binds to single-stranded DNA (ssDNA) and recruits over three dozen RPA-interacting proteins (RIPs) to coordinate multiple aspects of DNA metabolism including DNA replication, repair, and recombination. Rtt105 is a molecular chaperone that regulates nuclear localization of RPA. Whether and how Rtt105 regulates the activities of RPA is poorly understood. Here, we show that Rtt105 binds to multiple DNA binding and protein-interaction domains of RPA and configurationally staples the complex. In the absence of ssDNA, Rtt105 inhibits RPA binding to Rad52, thus preventing spurious binding to RPA-interacting proteins (RIPs). When ssDNA is available, Rtt105 promotes formation of high-density RPA nucleoprotein filaments and dissociates during this process. Free Rtt105 further stabilizes the RPA-ssDNA filaments by inhibiting RPA facilitated exchange. Collectively, our data suggest that Rtt105 sequesters free RPA in the nucleus to prevent untimely RIP interaction, while stabilizing RPA-ssDNA filaments at DNA lesion sites.

## Introduction

Replication protein A (RPA) is an essential single-stranded DNA (ssDNA) binding protein that coordinates almost all aspects of DNA metabolism including DNA replication, repair, and recombination (Wold, 1997). The myriad functions of RPA are facilitated through high affinity interactions with ssDNA (K_D_ < 10^−10^M) and physical interactions with over three dozen proteins (Fanning et al., 2006). With respect to the functions of RPA in double strand DNA break repair, current models posit that recognition of ssDNA by RPA triggers the DNA damage response (DDR). Coating of RPA on ssDNA serves as a nucleoprotein hub for the recruitment of several proteins including kinases that trigger downstream DNA repair pathways (Marechal and Zou, 2015). For example, yeast Mec1 or human ATM/ATR kinases are recruited on to the RPA-ssDNA substrate (Deshpande et al., 2017). How such protein-protein interactions with RPA are prevented in the absence of ssDNA remains a mystery.

Functional specificity for the myriad DNA metabolic functions of RPA is facilitated by a complex series of oligosaccharide/oligonucleotide binding (OB) folds/domains that are distributed across a heterotrimeric structural complex made of RPA70, RPA32 and RPA14 subunits (**Figure 1a**) (Bochkareva et al., 2002; Fan and Pavletich, 2012). OB-domains A, B, C, and F are in RPA70. OB-domain D and a winged-helix (wh) domain are situated in RPA32, and RPA14 houses OB-domain F. OB-F and the wh-domain are primarily responsible for mediating protein-protein interactions and are thus termed Protein-Interaction-Domains (PID^70N^ and PID^32C^, respectively). OB-A, B, C and D contribute most to ssDNA binding and are termed DNA-Binding-Domains (DBDs). The roles of OB-F and OB-E in binding ssDNA are uncertain. The DBDs and PIDs are connected by flexible linkers of varying lengths and thus can adopt a wide variety of structural assemblies on and off the DNA (Fanning et al., 2006), collectively termed ‘*configurational arrangements*’ in RPA. Conformational changes upon ssDNA binding and protein interactions are also quite extensive within the individual domains (Brosey et al., 2013).

**Figure 1.**
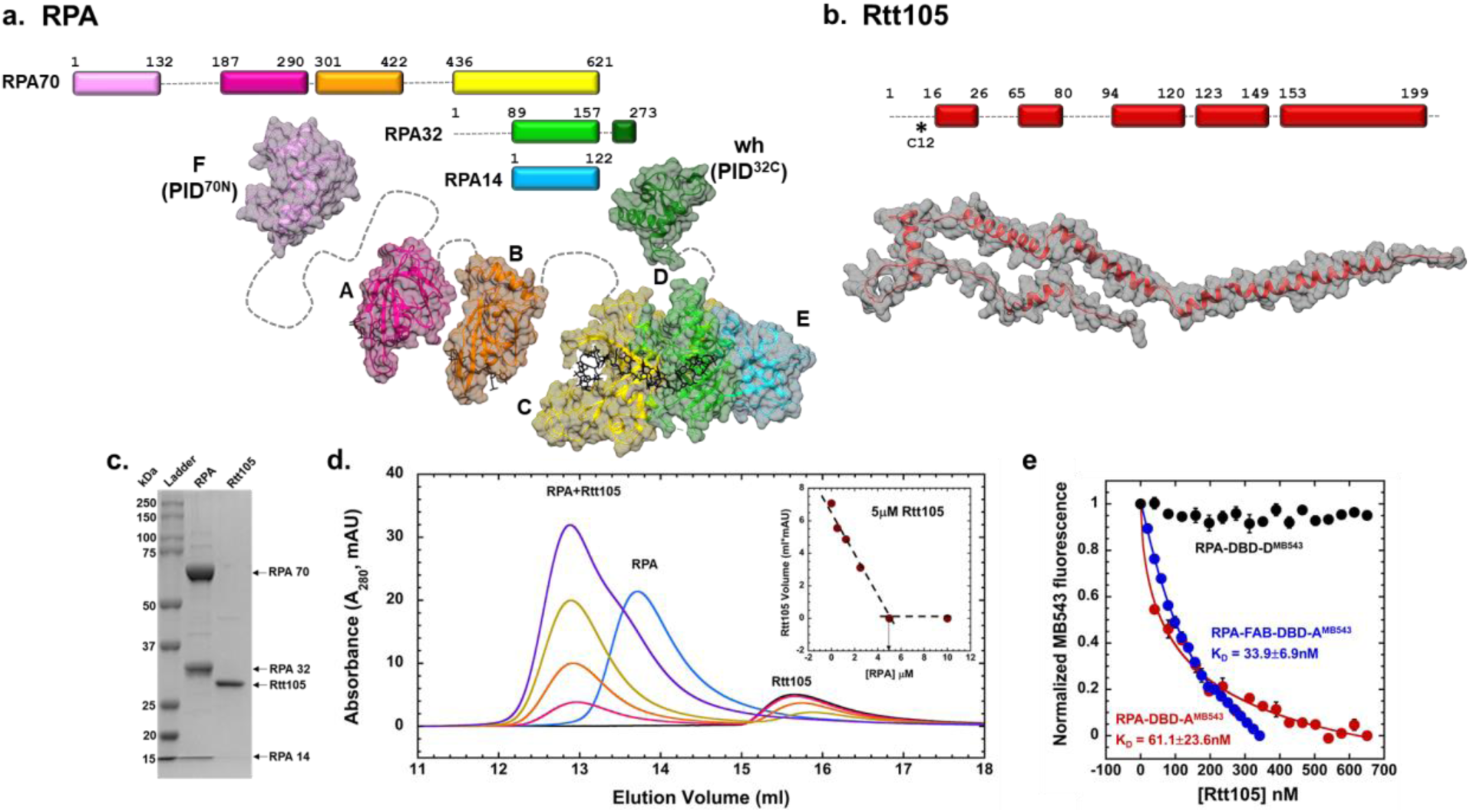
Rtt105 forms a stoichiometric complex with RPA. **a)** Schematic of the OB-domains of the three *S. cerevisiae* RPA subunits is depicted along with the structures of the individual domains. The domains are spread apart for clarity and the dotted lines denote the flexible linkers connecting the domains. DNA bound to the individual DBDs (A, B, C and D) is shown as sticks. **b)** A model of Rtt105 with predicted α-helices is shown (AlphaFold AF-P40063-F1). * denotes Cys-12 used to generate fluorescently labeled Rtt105. **c)** SDS-PAGE analysis of recombinantly purified RPA and Rtt105. **d)** Size exclusion chromatographic analysis of Rtt105 (5 µM) and RPA (0 – 5 µM) show RPA, Rtt105, and the Rtt105-RPA complex migrating as defined species. Insert shows the change in the Rtt105 peak volume as a function of RPA concentration and stoichiometric complex formation between Rtt105 and RPA. **e)** Changes in RPA-DBD-A^MB543^ or RPA-DBD-D^MB543^ fluorescence was measured after addition of increasing concentrations of Rtt105. Rtt105 binding does not influence the fluorescence of RPA-DBD-D^MB543^ but quenches RPA-DBD-A^MB543^ fluorescence. F-A-B labeled at DBD-A^MB543^ binding to Rtt105 reveals a higher binding affinity. Data were fit as described in the Methods to obtain the K_D_ values. SE from n=3 is shown.

In response to cellular DNA metabolic needs, RPA is shuttled from the cytoplasm into the nucleus. In *Saccharomyces cerevisiae*, the nuclear localization of RPA is facilitated by a chaperone-like protein called Regulator of Ty1 Transposition 105 (Rtt105) (Li et al., 2018). In higher eukaryotes, RPA-interacting protein (RPAIN or RIPα) appears to be the functional homolog. *RTT105* was originally discovered in a genome-wide screen as interacting with genes associated with genome maintenance pathways (Collins et al., 2007). Rtt105 physically interacts with RPA in the cytoplasm and mediates nuclear localization in complex with Kap95, a key karyopherin-beta protein that belongs to the importin-exportin family of nuclear transport protein complexes (Li et al., 2018). Consequently, deletion of Rtt105 or mutations that perturb the RPA-Rtt105 interaction give rise to defects in cell growth, DNA repair, and recombination (Corda et al., 2021; Wang et al., 2021).

Rtt105 is a 24 kDa protein (**Figure 1b**) and proposed to stabilize an extended conformation of RPA where all the OB-domains are stretched out (Li et al., 2019; Li et al., 2018; Wang et al., 2021). In addition to the nuclear transport functions, recent studies propose Rtt105 to enhance the ssDNA binding properties of RPA (Li et al., 2018; Wang et al., 2021). Depending on the number of DBDs bound to ssDNA, RPA can adopt distinct DNA binding modes where varying number of nucleotides are occluded by the RPA complex (Kumaran et al., 2006). The DBDs of RPA are also dynamic in nature and thus, while the protein is macroscopically bound to the ssDNA, each domain can exist in microscopically unbound states (Caldwell and Spies, 2020; Chen et al., 2016; Pokhrel et al., 2019). We recently showed that RPA can be envisioned to interact with ssDNA as two functional halves: a dynamic half consisting of the OB-F, DBD-A and DBD-B domains, and a less-dynamic half with DBD-C, DBD-D and OB-E (trimerization core or Tri-C) contributing more to the stability of RPA-ssDNA interactions (Ahmad et al., 2021).

RPA binds to ssDNA with very high affinity (K_D_ < 10^−10^M) (Kim et al., 1994; Pokhrel et al., 2019). Yet, a substantial increase in RPA binding affinity to ssDNA was shown in the presence of Rtt105 which led to a model where Rtt105 binds and promotes a configuration of RPA that extends out its DBDs such that maximal DNA binding contacts are promoted (Li et al., 2019; Li et al., 2018; Wang et al., 2021). Rtt105 inherently does not possess DNA binding activity and RPA intrinsically binds ssDNA with very high affinity. More strikingly, Rtt105 is not bound to the RPA-ssDNA complex. Thus, the finding that Rtt105 enhances the DNA binding activity of RPA, while not part of the DNA-RPA complex is perplexing, given that experimental tools such as electrophoretic mobility shift analysis (EMSA) do not have the required resolution to detect changes in sub-nanomolar biomolecular interactions. Furthermore, the rationale for Rtt105 stabilizing an extended conformation of RPA whilst having to transport it across the nuclear pore is also puzzling. To address these disparities, we undertook a detailed structural and mechanistic investigation of the Rtt105-RPA complex.

We here show that Rtt105 contacts multiple regions in RPA and conformationally compacts the multiple DBDs and PIDs through a configurational stapling mechanism. Shorter ssDNA substrates (∼15-35 nt) engage the Rtt105-RPA complex and promote remodeling of Rtt105 and a transition-state Rtt105-RPA-ssDNA complex is observed. However, Rtt105 does not change the affinity of RPA-ssDNA interactions. Longer ssDNA substrates (>35 nt) promote the binding of multiple RPA molecules and lead to dissociation of Rtt105. Interestingly, Rtt105 promotes binding of a higher density of RPA molecules which promote formation of RPA nucleoprotein filaments on DNA. In the absence of ssDNA, we find that Rtt105 blocks interactions between RPA and RPA-interacting proteins (RIPs) such as the homologous recombination mediator Rad52. ssDNA binding to Rtt105-RPA remodels the Rtt105-RPA-ssDNA complex and promotes RIP engagement. Posttranslational modifications of RPA further contribute to the remodeling of the RPA-Rtt105 complex. Finally, Rtt105 blocks the facilitated exchange activity of RPA thereby contributing to the stability of RPA nucleoprotein filaments. Thus, we here present a new functional role for Rtt105 where it sequesters RPA in the nucleus and serves as a negative regulator by blocking spurious interactions with RIPs in the absence of ssDNA.

## Results

### Rtt105 forms a stoichiometric complex with RPA

Rtt105 physically interacts with RPA and was shown to copurify as a complex (Li et al., 2018). To obtain the stoichiometry of the complex, using recombinantly purified proteins (**Figure 1c**) we analyzed formation of the Rtt105-RPA complex as a function of RPA concentration using size exclusion chromatography (SEC). We observe concentration-dependent stoichiometric binding between RPA and Rtt105 (**Figure 1d**). To determine a K_D_ value for the interaction, we followed the change in fluorescence in RPA upon complex formation with Rtt105. We used fluorescent versions of RPA where either DBD-A or DBD-D carry a site-specific environment-sensitive MB543 fluorophore (Kuppa et al., 2021; Pokhrel et al., 2019; Pokhrel et al., 2017). Upon binding to Rtt105 we observe a change in fluorescence for RPA-DBD-A^MB543^, but not for RPA-DBD-D^MB543^ (**Figure 1e**). Rtt105 binds to RPA with high affinity (K_D_=61.1±23.6 nM; **Figure 1e**) and the selective change in RPA-DBD-A^MB543^ fluorescence suggests that at least a part of Rtt105 is situated/bound in proximity to DBD-A. Furthermore, an F-A-B version of RPA containing just OB-F, DBD-A and DBD-D interacts with Rtt105 (**Supplemental Figure S1a**). Thus, the region around OB-F and DBD-A likely makes physical contacts with Rtt105. The F-A-B domain binds to Rtt105 with better affinity (K_D_=33.9±6.9 nM; **Figure 1e**). This data suggests that one or more of the Rtt105 binding sites in the F-A-B region is occluded in the context of the full-length RPA complex. A recent study suggested that a Val-106 to Ala substitution in OB-F (RPA70 subunit) was sufficient to abolish interaction with Rtt105 (Wang et al., 2021). However, in our analysis, this single point substitution in RPA (RPA^V106A^) is not sufficient to perturb the interaction (**Supplemental Figure S1b**) suggesting that additional contacts must exist between Rtt105 and RPA.

### Rtt105 does not enhance the DNA binding affinity of RPA, but alters the ssDNA bound configuration of DBDs

Recent studies have shown that Rtt105 enhances the ssDNA binding activity of RPA (Li et al., 2018). More perplexingly, Rtt105 is not bound to the RPA-ssDNA complex in pull-down experiments. Thus, how Rtt105 modulates the DNA binding activity of RPA, while not part of the RPA-ssDNA complex, is unresolved. In both these earlier studies, control experiments performed in the absence of Rtt105 showed only ∼50% of RPA bound to ssDNA oligonucleotides at equimolar concentrations (Li et al., 2018; Wang et al., 2021). These data are inconsistent with the highaffinity and stoichiometric DNA binding behavior of RPA (Pokhrel et al., 2019; Pokhrel et al., 2017). Thus, we first repeated these experiments, under similar experimental conditions, and do not observe this behavior. RPA binds stoichiometrically to a 5’-Cy5-(dT)_30_ oligonucleotide in electrophoretic mobility band-shift analysis (EMSA) and a preformed Rtt105-RPA complex does not influence the ssDNA binding activity of RPA (K_*D*_=5.5±1.1 and 6.7±2.1 nM for RPA and RPA-Rtt105, respectively; **Figures 2a & b**). Since EMSA experiments may not have the resolution to tease apart subtle differences in high-affinity biomolecular interactions, we used fluorescence anisotropy to further quantitate binding. RPA and the RPA-Rtt105 complex bound stoichiometrically to a 5’-FAM-(dT)_35_ ssDNA substrate and we do not see an Rtt105-induced enhancement of RPA-ssDNA binding activity (K_*D*_=4±1.3 and 2.4±1.2 nM for RPA and RPA-Rtt105, respectively; **Figure 2c**).

**Figure 2.**
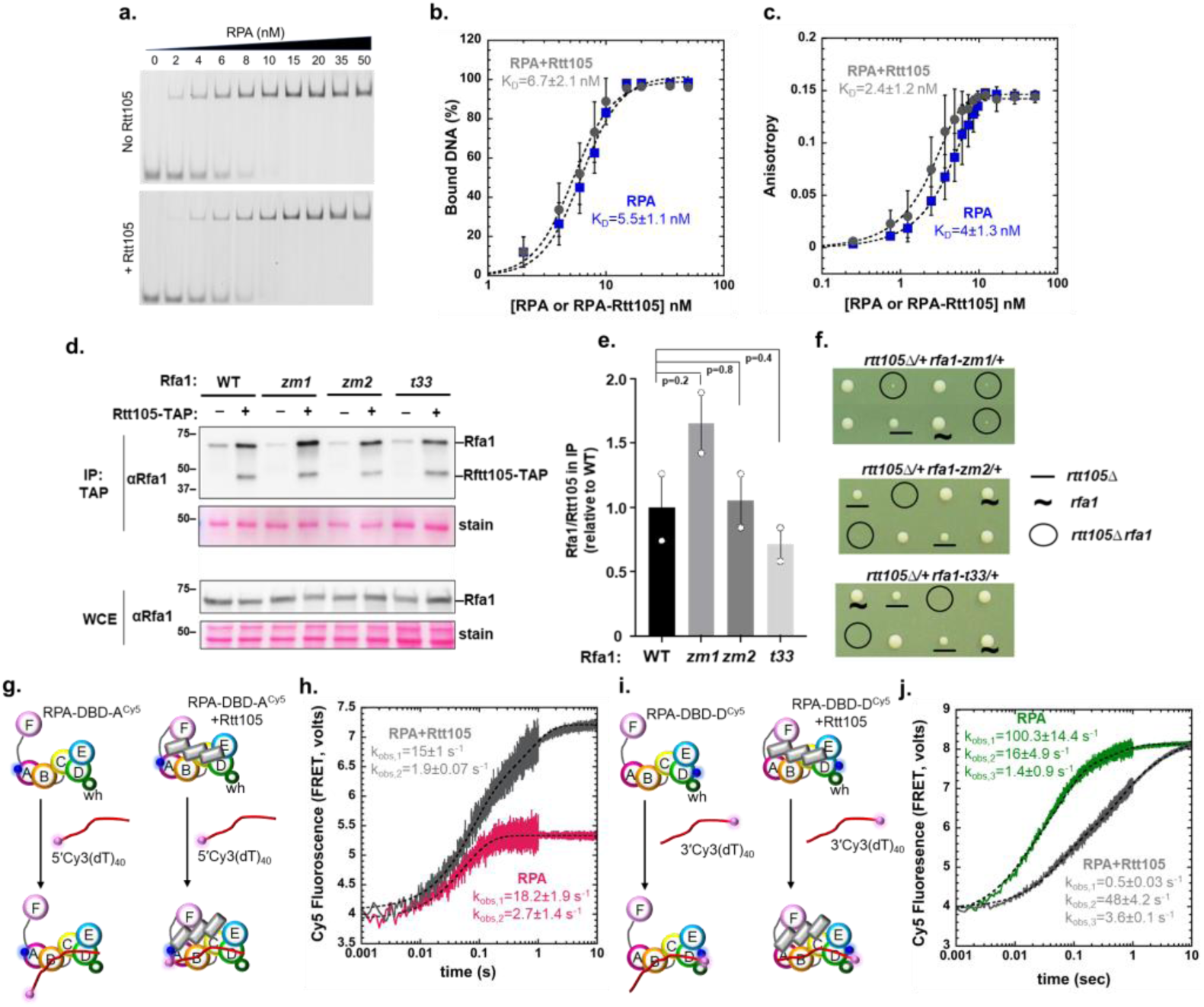
Rtt105 does not influence the DNA binding affinity of RPA but alters the configuration of DBD-A on ssDNA and kinetics of DBD-D binding. **a)** EMSA data showing RPA binding to 5′-Cy5-(dT)_30_ ssDNA in the presence or absence of Rtt105. **b)** Quantitation of EMSA gels in panel a show no appreciable difference in DNA binding affinity of RPA in the absence or presence of Rtt105. Binding data were fit as described in the Methods and yields K_D_=5.5±1.1 nM and 6.7±2.1 nM in the absence and presence of Rtt105, respectively. **c)** Fluorescence anisotropy measurements of RPA binding to 3′-FAM-(dT)_35_ measured in the absence or presence of Rtt105 shows no appreciable difference in DNA binding affinity. Data were fit as described in the Methods and yields K_D_=4±1.3 nM and 2.4±1.2 nM in the absence and presence of Rtt105, respectively. **d)** Rtt105 association with RPA is unaffected by its ssDNA binding ability. TAP-tagged Rtt105 was immunoprecipitated and RPA was detected with an anti-Rfa1 antibody. Representative immunoblots examining elute from the eluate (IP) and the whole cell extract (WCE) are shown. **e)** Graph represents the enrichment of Rfa1 present in the IP eluate relative to Rtt105 and normalized to wild type (WT). Mean and SEMs from two biological duplicates are plotted. P value obtained from Students t-test show that the differences in relative Rfa1 enrichment between WT Rfa1 and its mutants are not significant. **f)** *rtt105Δ* sensitizes *rfa1* mutants with impaired ssDNA association. Representative tetrads of diploids heterozygous for indicated mutations are shown. Spore clones were grown at 30 °C for 2 days. **g-h)** Stopped flow fluorescence measurements of DNA binding were monitored by FRET-induced enhancement of Cy5 fluorescence upon Cy3 excitation. FRET pairs positioned on the 5′-end of the DNA (5′-Cy3-(dT)_40_) and DBD-A of RPA (RPA-DBD-A^Cy5^) show similar rates of DNA-induced changes in the absence and presence of Rtt105 (k_obs,1_=18.2±1.9 s^-1^, 15±1 s^-1^; k_obs,2_=2.7±1.4 s^-1^, 1.9±0.07 s^-1^; in the absence and presence of Rtt105, respectively), but Rtt105 markedly enhances the amplitude of Cy5 fluorescence. Data were fit to a double exponential equation as described in the Methods. **j-k)** Similar experiments performed with FRET pairs positioned on the 3′- end of the DNA (3′-Cy3-(dT)_40_) and DBD-D of RPA (RPA-DBD-D^Cy5^) show an Rtt105 induced reduction in the rate of DNA binding (k_obs,1_=100.3±14.4 s^-1^, 0.5±0.03s^-1^; k_obs,2_=16±4.9, 48±4.2 s^-1^; k_obs,3_=1.4±0.9, 3.6±0.1 s^-1^in the absence and presence of Rtt105, respectively). But, unlike in panel d, both traces show similar changes in fluorescence amplitude. Data were fit to a triple exponential equation as described in the Methods. SE from n=3 independent experiments are shown.

These results suggest that to a large degree, RPA association with Rtt105 is a biochemical feature that can be functionally distinct from its ssDNA binding activity. This notion of two separable features of RPA is supported by our *in vivo* data, where we tested the cellular effects of deletion of Rtt105 along with three distinct DNA binding mutants of RPA. zm1 (K494A) and zm2 (N492D, K494R, K494R) mutants of RPA carry amino acid substitutions close to the Zn^2+^-finger binding domains in DBD-C and have reduced ssDNA binding activity (Dhingra et al., 2021). t33 is a well characterized RPA mutant with a S373P substitution in DBD-B (Chen et al., 1998; Deng et al., 2014). Immunoprecipitation of Rtt105 from cells expressing these mutant variants of RPA show that the physical interaction between the two proteins is not perturbed by these mutations in RPA (**Figures 2d & e**). From a genetic point view, the lack of influence of RPA DNA binding by Rtt105 (**Figures 2b & c**) predicts that mutants impairing RPA binding to ssDNA in cells lacking Rtt105 should have additive genetic interactions. Indeed, we find that RPA mutations that displayed reduced ssDNA binding activities were additive when combined with *rtt105* null cells when assayed for growth (**Figure 2f**). These *in vivo* data support the idea that RPA-Rtt105 interactions and RPA-ssDNA binding are two separable functions.

### Rtt105 mildly influences the ssDNA binding kinetics and configurations of individual DBDs of RPA

Since Rtt105 does not change the equilibrium ssDNA binding properties of RPA, we next tested whether the kinetics of RPA-ssDNA interactions were influenced by Rtt105. We used fluorescent RPA carrying Cy5 on either DBD-A or DBD-D and captured the kinetics of binding to (dT)_40_ oligonucleotides labeled with Cy3 at either the 5’ or 3’ end. Changes in Förster resonance energy transfer (FRET) induced Cy5 fluorescence were monitored by exciting Cy3 in a stopped flow fluorometer (**Figures 2g-j**). Since the DBDs of RPA bind ssDNA with defined polarity (Iftode and Borowiec, 2000; Kolpashchikov et al., 2001), with DBD-A and DBD-D residing close to the 5’and 3’ ends, respectively, the change in fluorescence in each experiment reflects the configurational changes of the respective DBDs with respect to their cognate DNA termini positions (Pokhrel et al., 2019). The data show that DBD-A binds with ∼similar rates to the 5’ endlabeled DNA in the absence or presence of Rtt105 (**Figures 2g & h**). However, the fluorescence signal amplitude in the presence of Rtt105 is twice that observed for RPA alone (**Figure 2h**). In contrast, Rtt105 reduces the rate of DBD-D binding to the 3’ end-labeled DNA (**Figures 2i & j**), but the signal amplitudes are similar (**Figure 2j**). These data suggest that Rtt105 differentially influences the two DBDs of RPA but does not inform whether Rtt105 dissociates from RPA. In addition, these findings supports a model where Rtt105 influences the configuration of the domains of RPA, but not the affinity of the the RPA-ssDNA interaction.

### Rtt105-RPA-ssDNA complexes are detected on short DNA substrates

To directly test formation of the Rtt105-RPA-ssDNA complex we performed SEC analysis of the individual proteins and their complexes. Rtt105 (24 kDa) and RPA (114 kDa) migrate as distinct species due to large differences in their molecular weight and migrate together as a complex when pre-mixed (**Figure 3a**). However, when equimolar amounts of (dT)_35_ are added to the complex, Rtt105 primarily remains bound as a higher order Rtt105-RPA-(dT)_35_ complex. A small fraction of free Rtt105 and RPA-(dT)_35_ complexes are also observed (**Figure 3a & Supplemental Figure S1c-d**). These data show that the configurational changes in RPA observed in the stopped flow experiments (**Figure 2**) are induced by both Rtt105 and DNA. We propose that the RPA-Rtt105 complex binds ssDNA and both proteins are likely reconfigured as the DBDs engage onto ssDNA. Thus, the interactions between RPA and Rtt105 are different in the DNA bound versus unbound states. Based on the changes in observed fluorescence between the ends of the DNA and the DBDs, and the higher Rtt105 binding affinity for the F-A-B half of RPA, we propose that upon ssDNA binding Rtt105 interactions with RPA are likely shifted towards the F-A-B part of RPA. This model suggests an active repositioning of Rtt105 on RPA upon ssDNA binding.

**Figure 3.**
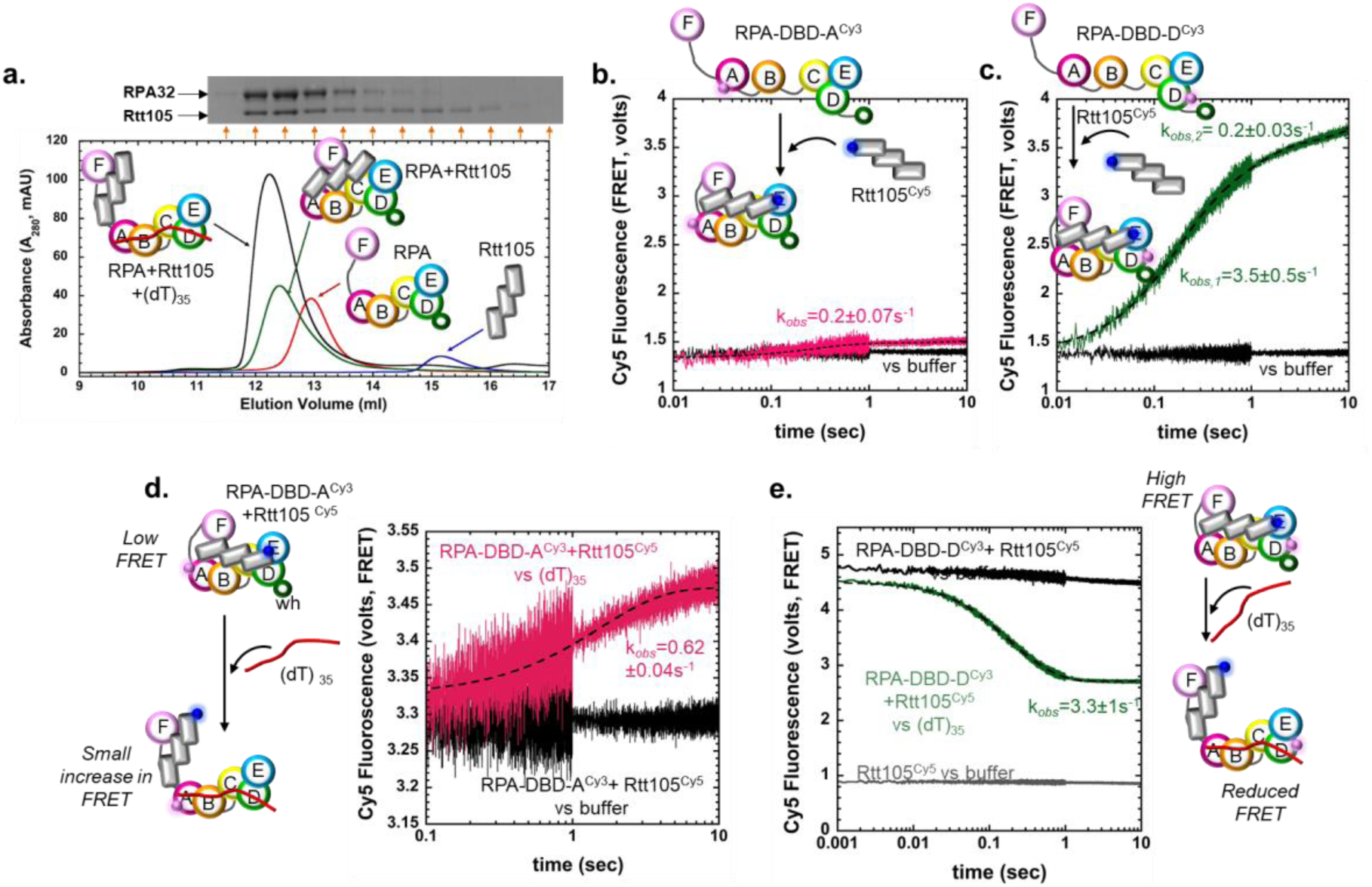
Rtt105 forms a higher order complex with RPA bound to short ssDNA substrates. **a)** Size exclusion chromatography (SEC) analysis of Rtt105, RPA, and Rtt105-RPA show complex formation between the proteins. Addition of equimolar amounts of (dT)_35_ drive formation of a higher order Rtt105-RPA-(dT)_35_ complex. SDS page analysis of the SEC fractions (insert) shows RPA and Rtt105 comigrating as a complex in the presence of DNA. The kinetics of RPA binding to Rtt105 was measured using FRET with Cy5-labeled Rtt105 and either **b)** RPA-DBD-A^Cy3^ or **c)** RPA-DBD-D^Cy3^. Cy3 positioned on DBD-D produces a larger FRET-induced Cy5 enhancement compared to DBD-A suggesting that the N-terminal region of Rtt105 resides closer to DBD-D of RPA. The data for RPA-DBD-A in panel b fits well to a single exponential model (k_obs_=0.62±0.73 s^-1^) whereas the RPA-DBD-D data is better described by a double exponential (k_obs,1_=3.5±0.5s^-1^, k_obs,2_=0.2±0.03s^-1^). These data suggest faster, initial engagement of Rtt105 towards the trimerization core (Tri-C) of RPA. Stopped flow experiments show transition from either a preformed **d)** RPA-DBD-A^Cy3^:Rtt105^Cy5^ or **e)** RPA-DBD-D^Cy3^:Rtt105^Cy5^ complex to remodeled Rtt105^Cy5^-RPA^Cy3^-(dT)_35_complexes upon addition of ssDNA. A small increase in Cy5 fluorescence is observed for the RPA-DBD-A^Cy3^:Rtt105^Cy5^ complex. In contrast, a sharp decrease in Cy5 signal is observed for the RPA-DBD-D^Cy3^:Rtt105^Cy5^ complex. The Cy5 fluorescence does not reach the baseline signal for Rtt105^Cy5^. The remodeling data fits well to a single step model (k_obs_=3.3±0.96 s^-1^). Representative stopped flow data averaged from seven to eight shots from one experiment are shown. SD was calculated from three independent experiments (n=3).

To experimentally capture this process, we generated fluorescently labeled Rtt105 to perform FRET analysis of the RPA-Rtt105 complex. Rtt105 has two non-conserved native Cys residues situated adjacently (Cys12 and Cys13, **Supplemental Figure S2a**). Substitution of either or both Cys residues to Ser does not change the secondary structure, RPA binding, or RPA remodeling activity of Rtt105 (**Supplemental Figure S2**). Thus, we generated a single Cys version of Rtt105 by converting Cys13 to Ser and fluorescently labeled Rtt105^C13S^ with Cy5 using maleimide chemistry (**Supplemental Figures S2e-f**). Generation of a functional fluorescent Rtt105 enabled us to investigate the assembly, remodeling, and disassembly of the Rtt105-RPA complex using stopped flow FRET. First, we measured the kinetics of Rtt105^Cy5^ binding to RPA-DBD-A^Cy3^ or RPA-DBD-D^Cy3^ by exciting Cy3 and monitoring the changes in Cy5 fluorescence. A small increase in FRET-induced Cy5 fluorescence is observed when Rtt105^Cy5^ binds to RPA-DBD-A^Cy3^ (**Figure 3b**), but a robust signal change is observed for RPA-DBD-D^Cy3^ (**Figure 3c**). These data suggest that in the absence of DNA, the N-terminal region of Rtt105 (where the Cy5 is positioned) is situated closer to DBD-D. The data for DBD-A^Cy3^ and Rtt105^Cy5^ binding fits to a single-step model (k_*obs*_=0.2±0.07s^-1^; **Figure 3b**) whereas the DBD-D^Cy3^ and Rtt105^Cy5^ data fits better to a twostep model (k_*obs,1*_=3.5±0.5s^-1^, k_*obs,2*_=0.2±0.03s^-1^; **Figure 3c**). Conservative interpretation of the kinetics suggests faster initial binding of the N-terminal region of Rtt105 closer to DBD-D followed by slower binding/rearrangements to other regions of RPA.

Next, to obtain the kinetics of ssDNA-induced remodeling of the Rtt105-RPA complex, we preformed either the low-FRET RPA-DBD-A^Cy3^:Rtt105^Cy5^ or high-FRET RPA-DBD-D^Cy3^:Rtt105^Cy5^ or complex and monitored the change in Cy5 fluorescence upon addition of (dT)_35_ ssDNA. When the preformed RPA-DBD-A^Cy3^:Rtt105^Cy5^ complex is mixed with (dT)_35_, a small enhancement in Cy5 fluorescence is observed (k_*obs*_=0.62±0.04 s^-1^, **Figure 3d**). In the corollary experiment, a rapid, single-step transition from the high-FRET to low-FRET state is observed when the RPA-DBD-D^Cy3^:Rtt105^Cy5^ complex is mixed with (dT)_35_ (k_*obs*_=3.3±1 s^-1^; **Figure 3e**). However, the fluorescence signal does not decrease to the baseline Rtt105^Cy5^ level. These data further support a model where Rtt105 does not completely dissociate from the RPA-(dT)_35_ complex but is remodeled such that the N-terminal region of Rtt105 is moved away from DBD-D upon DNA binding. This movement likely encompasses movement of Rtt105 from the trimerization core (DBD-C, RPA32, and RPA14; Tri-C) and towards the F-A-B region (OB-F, DBD-A and DBD-B) of RPA. Since ssDNA binding to RPA causes significant configurational changes, we next tested if free Rtt105 in solution can bind to preformed RPA-ssDNA complexes. FRET pairs were positioned on RPA (either RPA-DBD-A^Cy3^ or RPA-DBD-D^Cy3^) and Rtt105 (Rtt105^Cy5^). Stoichiometrically bound RPA^Cy3^-(dT)_35_ complexes were preformed and mixed with equimolar amounts of Rtt105^Cy5^. In both cases, only a very small change in Cy5 fluorescence is observed (**Supplemental Figures S3a & b**) suggesting that Rtt105 poorly binds to pre-formed RPA-DNA complexes. We do note that when RPA is in complex with ssDNA, its F-A-B is likely more exposed (described below) and thus, within the small fraction where Rtt105 binding is observed, we see a faster rate of Rtt105 binding to the F-A-B region (**Supplemental Figures S3c**).

### Rtt105 configurationally staples RPA through contacts with multiple domains in RPA

A structure of the Rtt105-RPA complex is not available and our efforts to obtain one using cryo-electron microscopy have not been successful because of the dynamic nature of the interactions (data not shown). Thus, to understand how Rtt105 interacts with RPA and decipher how the complex is remodeled by ssDNA, we performed cross-linking mass spectrometry (XL-MS) and hydrogen-deuterium exchange mass spectrometry (HDX-MS) analysis of the Rtt105-RPA complex in the absence or presence of DNA. We observe excellent peptide coverage for both RPA and Rtt105 in MS analysis (92% RPA70, 67% RPA32, 74% RPA14, and 67% Rtt105; **Supplemental Figure S4**) and thus can comprehensively assess the global conformational changes.

For XL-MS, RPA or RPA-Rtt105 were cross-linked with bis(sulphosuccinimidyl)suberate (BS3), which reacts with primary amines in lysine side chains and the N-termini (Rappsilber, 2011). RPA and Rtt105 cross-linked readily, as observed by the shift in the protein bands in SDS-PAGE analysis after BS3 addition (**Supplemental Figure S5**). MS analysis of the crosslinked peptides yielded several linkage pairs for the RPA and RPA-Rtt105 complexes, respectively (**Figure 4a-e**). The data mapped onto the individual subunits of RPA reveal extensive crosslinks between the N-terminal two-thirds of Rtt105 and all DBDs and PIDs in RPA (**Figures 4a-c**). Multiple contacts to OB-F, DBD-A, DBD-B, and DBD-C (all in RPA70) are observed. The F-A and A-B flexible linkers show no crosslinks, but a few are observed in the B-C linker (**Figure 4a**). A limited number of crosslinks are observed in RPA32 with one peptide in DBD-D making 9 contacts in Rtt105 (**Figure 4b**). Finally, a single crosslink is observed in OB-E (RPA14; **Figure 4c**). These data agree with our observation that shows higher affinity interactions between Rtt105 and the F-A-B half of RPA (**Figure 1e**).

**Figure 4.**
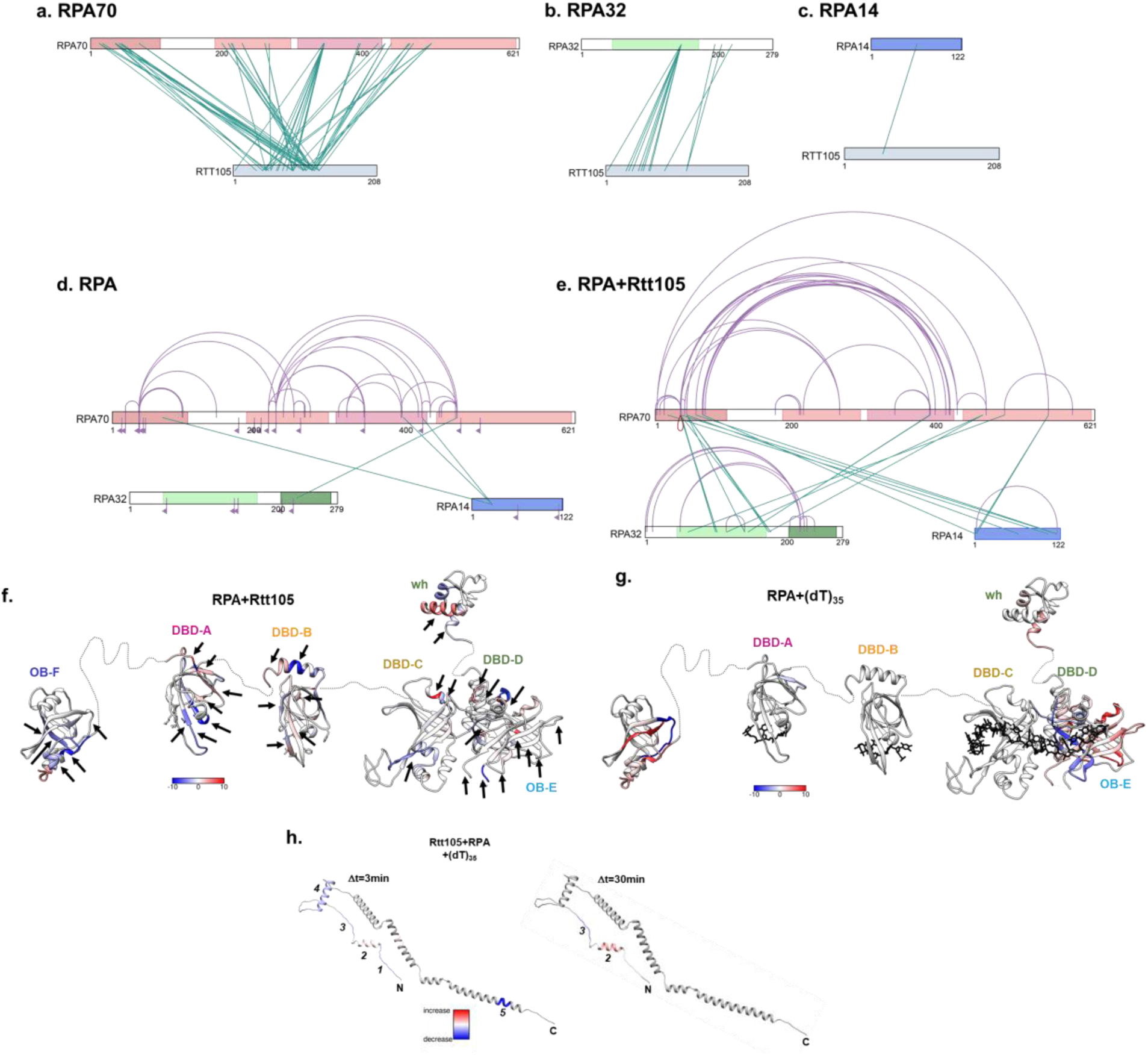
Rtt105 configurationally staples RPA through multiple contacts with DBDs and PIDs. Crosslinking mass spectrometry (XL-MS) analysis of the RPA-Rtt105 complex reveals crosslinks between Rtt105 and **a)** RPA70, **b)** RPA32, and **c)** RPA14 subunits. Crosslinks are observed in all OB-folds and the BC-linker. Intersubunit (green lines) and intra-subunit (purple traces) crosslinks in RPA are shown in the **e)** absence or **f)** presence of Rtt105, respectively. An increase in both inter-and intra-subunit crosslinks are observed upon Rtt105 binding to RPA. The inverted flag marks in panel d denote the sites of mono-crosslinks and show that they have no partner sites available in proximity for the BS3 crosslinker due to the extended configuration of RPA in the absence of Rtt105. Hydrogen-deuterium mass spectrometry (HDX-MS) analysis of **f)** RPA-Rtt105 and **g)** RPA-(dT)_35_ complexes show net changes in deuterium uptake or loss (ΔHDX) in almost all domains of RPA. The arrows point to ΔHDX that are unique to the Rtt105-RPA complex and not observed in the RPA-(dT)_35_ complex. **h)** ΔHDX changes in Rtt105 bound to RPA in the absence and presence of (dT)_35_ are denoted in the AlphaFold derived model of Rtt105. The regions of change are denoted 1-5. The ΔHDX changes upon DNA binding suggests remodeling of Rtt105. The scales (blue to red) denote the net ΔHDX.

A closer look at the crosslinked peptides in the individual DBDs reveal that the crosslinks in DBD-A and DBD-B are not in the DNA binding pockets (**Supplemental Figures S6a & b**). In contrast, some of the crosslinked peptides in DBD-C and DBD-D are in the DNA binding region (**Supplemental Figures S6c & d**). Based on the predicted structure and charge distribution of Rtt105 (**Supplemental Figure S6e**), the elongated helical regions likely extend and bind across the many domains of RPA. Circular dichroism (CD) analysis of Rtt105 confirm the helical nature of Rtt105 (**Supplemental Figure S2b**). Due to the extensive crosslinks observed we propose a *configurational stapling* model for the RPA-Rtt105 interaction. In this model, Rtt105 can be envisioned sitting on the surface of RPA, interacting with multiple domains, and configurationally constraining (or stapling) the DBDs and PIDs that are connected by flexible linkers.

Comparison of XL-MS data for RPA in the absence or presence of Rtt105 lends further support to the configurational stapling model. In the absence of Rtt105, a total of four inter-subunits crosslinks are captured between the three RPA subunits (**Figure 4d**). In contract, Rtt105 binding induces 16 distinct inter-subunits crosslinks (**Figure 4e**). These data show that the three subunits are brought in proximity upon Rtt105 binding. Analysis of the intra-subunit crosslinks further highlight the configurational stapling mechanism as the crosslinks between the individual DBDs and PIDs (within each subunit) are extensively increased in the presence of Rtt105 (**Figures 4d & e**). Thus, in contrast to existing models that propose a stretching of the RPA domains upon Rtt105 binding (Li et al., 2019; Li et al., 2018; Wang et al., 2021), we show that the many DBDs and PIDs in RPA are compacted by Rtt105 through a configurational stapling mechanism.

To further assess the degree of configurational and conformational changes in RPA induced by Rtt105, we performed HDX-MS analysis of RPA in the absence and presence of Rtt105 (**Figures 4f-g** and **Supplemental Figures S7-S11**). In agreement with the XL-MS observations, changes in deuterium uptake/loss are observed in all the DBDs and PIDs of RPA. Most of the changes favor an increase in deuterium uptake (denoted as red) suggesting enhanced solvent exposure of RPA domains upon Rtt105 binding. To delineate whether Rtt105 binding overlaps with ssDNA binding to RPA, we compared the ΔHDX between the Rtt105-RPA and RPA-ssDNA complexes (**Figures 4f & g**; regions marked with arrows are unique to Rtt105 binding). Thus, ssDNA ((dT)_35_) binding remodels the Rtt105-RPA complex in agreement with the kinetic data.

Additional support for the remodeling of the RPA-Rtt105 complex by ssDNA is evident in the ΔHDX changes observed within Rtt105. Upon addition of DNA to the preformed Rtt105-RPA complex we observe deuterium uptake/release in five distinct regions, primarily around the N-terminal half (**Figure 4h** and **Supplemental Figure S12**). When assessed as a function of time, the remodeling of the N-terminal region of Rtt105 is observed at shorter time points. At later time points, the exchange patterns are preserved in only two of the original five regions. These data further support our model where ssDNA binding remodels the N-terminal region of Rtt105 away from Tri-C region of RPA and Rtt105 interactions with the RPA-ssDNA complex are maintained through interactions between Rtt105 and F-A-B region of RPA. The precise nature of the interactions will have to be established through structural studies.

### Length of ssDNA and assembly of multiple RPA molecules promote release of Rtt105

In pull-down and single-molecule DNA curtain analysis Rtt105 has been proposed to not remain in complex with RPA on DNA (Li et al., 2019; Li et al., 2018; Wang et al., 2021). Since we clearly observe a Rtt105-RPA-ssDNA complex on short DNA oligonucleotides (e.g. (dT)_35_, **Figure 3a**) we next tested whether DNA length influenced Rtt105 dissociation. Longer ssDNA substrates offer binding sites for more than one RPA molecule. In this scenario, we previously showed that DBD-A from one RPA can interact with DBD-E of the neighboring RPA while being stably bound to DNA through Tri-C-DNA interactions (Yates et al., 2018). We hypothesized that since the F-A-B region is in an alternate configuration when multiple RPA are bound on DNA, Rtt105 might fully dissociate from the RPA-ssDNA complex. Using SEC, we tested whether Rtt105 remained bound to RPA when (dT)_70_ ssDNA was added to the reaction (**Figure 5a**). The elution profile shows three distinct species: [RPA]_2_-(dT)_70_, Rtt105-RPA-(dT)_70_, and free Rtt105. The larger, earlier eluting, species does not contain Rtt105 (**Figure 5a** and **Supplemental Figure S13**). Thus, assembly of multiple RPA molecules serves as a trigger for Rtt105 dissociation.

**Figure 5.**
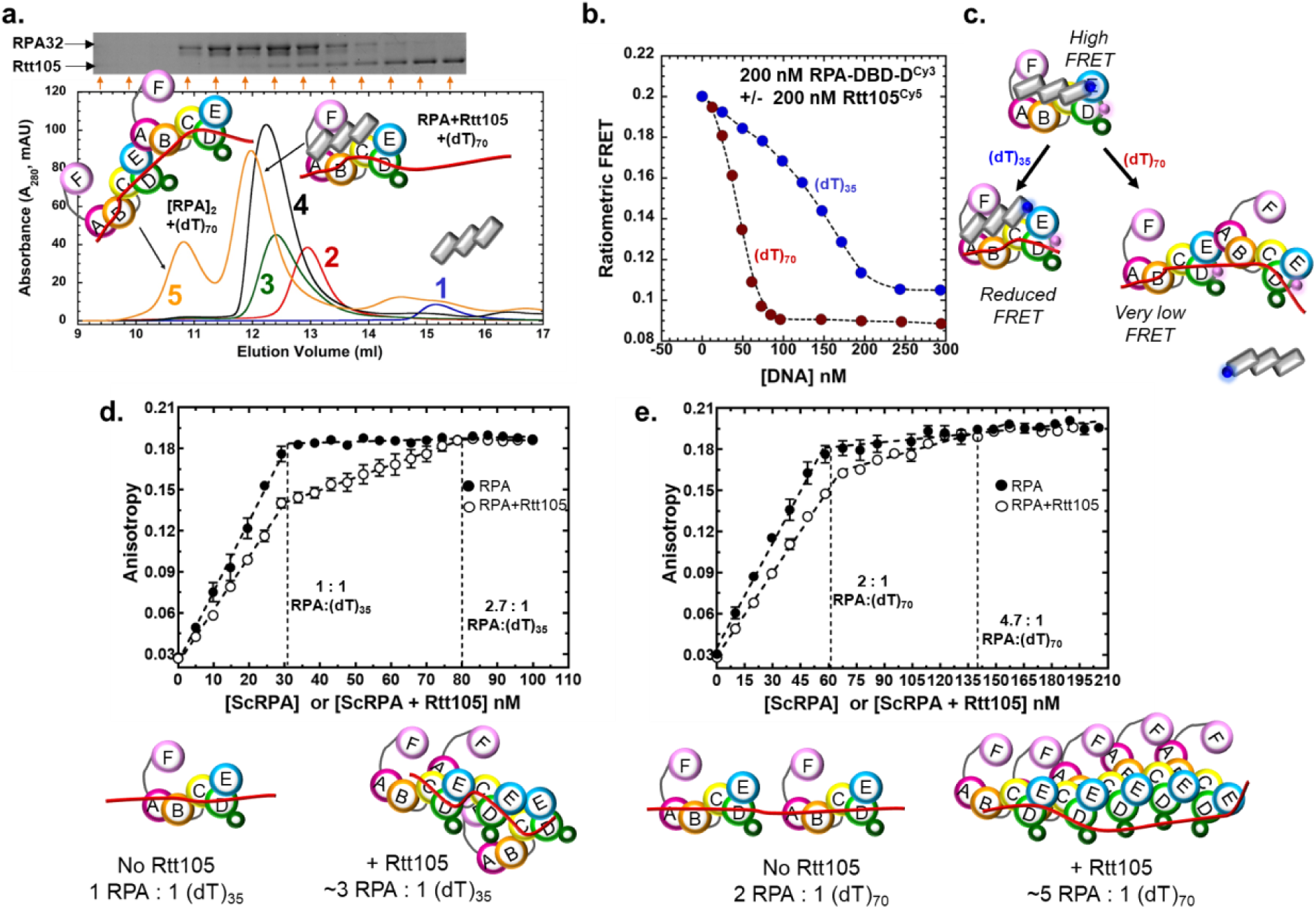
Binding of multiple RPA molecules on longer ssDNA substrates triggers the release of Rtt105 and formation of a high-density RPA nucleoprotein filament. **a)** Size exclusion chromatography (SEC) analysis of the Rtt105-RPA complex in the presence of equimolar amounts of (dT)_70_ show the presence of three distinct species. A larger [RPA]_2_-(dT)_70_] and smaller remodeled [Rtt105-RPA-(dT)_70_]^*^ complex are observed. Rtt105 is displaced from the larger complex and seen as the third peak. The distribution of Rtt105 in the three peaks can be noted in the SDS-PAGE gel shown as insert. Traces 1, 2, 3 and 4 are shown as reference and denote Rtt105, RPA, Rtt105-RPA, and the Rtt105-RPA-(dT)_35_ complex, respectively. **b)** Preformed RPA-DBD-D^Cy3^-Rtt105^Cy5^ complexes were mixed with increasing concentrations of either (dT)_35_ or (dT)_70_ and the change in FRET was measured. At ∼2:1 ratios of RPA:(dT)_70_ the FRET signal is significantly reduced. At ∼1:1 ratio and lower, an alternate reduced FRET state is observed for (dT)_35_. **c)** The reduced FRET state likely corresponds to the remodeled [Rtt105-RPA-(dT)_35_]^*^ complex and the very low FRET state corresponds to the [[RPA]_2_-(dT)_70_] state lacking Rtt105. **d)** Fluorescence anisotropy experiments were performed with 30 nM (dT)_35_ ssDNA by adding increasing amounts of RPA or the RPA:Rtt105 complex. On (dT)_35_, RPA binds with 1:1 stoichiometry, but the Rtt105 increases the binding density of RPA to ∼3 RPA/(dT)_35_ molecule. **e)** A similar phenomenon is observed on the longer (dT)_70_ substrate where a higher density of RPA binding (∼5 RPA/(dT)_70_) is observed. SE from three independent experiments are shown.

To further probe the relationship between DNA length, multiple RPA binding, and Rtt105 dissociation, we used the FRET signal between RPA-DBD-D^Cy3^ and Rtt105^Cy5^ and followed complex dynamics as a function of either (dT)_35_ or (dT)_70_ concentration (**Figure 5b**). Mixing 200 nM each of RPA-DBD-D^Cy3^ and Rtt105 ^Cy5^ yields a high FRET state, and the signal decreases as DNA is added to the reaction. (dT)_70_ drives dissociation of Rtt105 at lower DNA concentrations (∼100 nM) and produces a very low FRET state. At these ratios (2:1, RPA:(dT)_70_) multiple RPA are expected to be bound to a single DNA molecule. In contrast, similar experiments done with (dT)_35_ results in formation of an alternate low-FRET [Rtt105:RPA:(dT)_35_]^*^ state that corresponds to the remodeled complex where most Rtt105 remains bound, but in a remodeled state. Furthermore, the longer (dT)_70_ substrate is 3-fold better at arriving at the low FRET state compared to (dT)_35_ (K_1/2_=130.4±10.5 and 42.8±0.37 nM for (dT)_35_ and (dT)_70_, respectively; **Figure 5b**). Performing this experiment as a function of ssDNA length reveals that ssDNA ∼35 nt and longer are effective in promoting Rtt105 dissociation from RPA (**Supplemental Figure S14**). Thus, when adequate ssDNA is present to saturate all the individual DBDs (≤ (dT)_35_) Rtt105 is remodeled on RPA. Longer DNA (≥(dT)_35_), and higher RPA:DNA ratios promote binding of multiple RPA molecules and drive complete Rtt105 dissociation (**Figure 5c**).

### Rtt105 drives formation of high-density RPA-ssDNA filaments

In single molecule DNA curtain experiments, Rtt105 was shown to drive formation of stretched RPA-ssDNA filaments (Li et al., 2018). Since Rtt105 was proposed to stretch out the domains of RPA, the model invoked elongated RPA bound to longer ssDNA with Rtt105 not bound (Li et al., 2019). Our data show a compaction of RPA by Rtt105. To reconcile these differences, we wondered if Rtt105 promoted alternate DNA-bound RPA configurations. For example, if Rtt105 were bound to the F-A-B part of RPA, one could envision an RPA-ssDNA filament where multiple RPA are bound using only the Tri-C region. Since the Tri-C region is more stably bound to ssDNA (Ahmad et al., 2021), the resulting filament could be more rigid and elongated as observed in the DNA curtain experiments. To test this model, we used fluorescence anisotropy to monitor RPA binding to (dT)_35_ or (dT)_70_ ssDNA in the absence or presence of Rtt105. On the shorter (dT)_35_ substrate, the signal saturates at 1:1 (RPA:DNA) in the absence of Rtt105 (**Figure 5d**). In contrast, the ratio shifts to 2.7:1 (RPA:DNA) when the RPA-Rtt105 complex is used (**Figure 5d**). On the longer (dT)_70_ substrate, this effect is further exaggerated as ∼5 molecules of RPA are loaded onto the DNA in the presence of Rtt105 compared to ∼2 when no Rtt105 is present in the reaction (**Figure 5e**). Thus, remarkably, the configuration of RPA on DNA formed in the presence of Rtt105 is stable and distinctly different with a higher density of RPA molecules bound to the DNA. Interestingly, the difference in the amplitude of the anisotropy signals at the 1:1 fluctuation point for both DNA substrates suggest that the hydrodynamic radius of the RPA-DNA complex in the absence of Rtt105 is larger. We propose that the RPA molecules formed in the presence of Rtt105 likely have the F-A-B region (the dynamic half of RPA) not bound to DNA.

In all the above experiments, we investigated modulation of the RPA-Rtt105 complex using short ssDNA oligonucleotides. To test whether our observations of RPA-Rtt105 remodeling were recapitulated on longer kilobase-long ssDNA we utilized C-trap analysis. Here lambda DNA (∼48.5 kbp) is tethered between two beads and stretched to produce ssDNA (**Supplemental Figures S15a-c**). Fluorescent proteins binding to DNA, remodeling, and/or dissociating are visualized as kymographs as a function of time. Rtt105^Cy5^ alone does not interact with DNA at either low or high concentrations and thus no fluorescent spots are observed (**Supplemental Figure S15d**). When the Rtt105^Cy5^-RPA complex is premixed at 1:1 ratio and first incubated with the DNA in the first chamber and then moved to chamber containing buffer, a few spots of Rtt105^Cy5^ are occasionally encountered (**Supplemental Figures S15e-f**). Even in these rare instances, Rtt105^Cy5^ dissociates from these spots with a t_1/2_∼5s. Quantitation of these data over multiple experiments reveal a distribution of 1-4 binding/dissociation events per kymograph recorded (**Supplemental Figure S15f**). These experiments provide direct visual evidence for engagement of Rtt105-RPA onto DNA and Rtt105 dissociation under conditions where multiple RPA can bind onto a single DNA lattice and form a nucleoprotein filament.

### Rtt105 impedes facilitated exchange of RPA

Free RPA has been shown to exchange with RPA bound to ssDNA through a process called facilitated exchange (FE) (Ma et al., 2017). This activity arises from the dynamic binding, rearrangement, and dissociation of one or more DBDs on the DNA and thus transiently exposed short segments of ssDNA allow free RPA to gain access. In this scenario, while multiple RPA molecules are bound on DNA, we expect only short segments DNA exposed at any given point during FE. The biological significance of FE is poorly understood but thought to contribute to RPA dynamics and RPA-promoted enhancement or impediment of DNA metabolic processes in the cell. FE could also be envisioned as a counterproductive process in the cell when excess RPA is around as the stability of a RPA nucleoprotein filament will be perturbed. We hypothesized that Rtt105 might influence FE by serving as a sink to sequester free RPA. To test this idea, we performed FE experiments using RPA labeled with FRET pairs. Binding of equimolar amounts of RPA-DBD-A^Cy3^ and RPA-DBD-D^Cy5^ (100 nM each) to (dT)_97_ ssDNA (30 nM) results in a high FRET signal as multiple RPA molecules assemble together (**Figure 6a**). When excess unlabeled RPA is added to a preformed RPA-DBD-A^Cy3^:RPA-DBD-D^Cy5^:(dT)_97_ complex, FE occurs leading to a loss in the FRET signal (**Figure 6b**, red trace). In contrast, when challenged with RPA prebound to Rtt105, no FE is observed, and the FRET signal is not perturbed (**Figure 6b**, blue trace). Thus, the short segments of free ssDNA available during FE does not allow binding of the Rtt105-RPA complex. To further investigate whether Rtt105 blocks FE on long RPA nucleoprotein filaments, we used C-trap analysis where lambda DNA (∼48.5 kbp) was first coated with fluorescent RPA (RPA-DBD-D^MB543^) and then moved over to a channel containing either buffer in absence or presence of unlabeled RPA or RPA+Rtt105 (**Figure 6c-e**). Extensive FE is observed when unlabeled RPA is introduced as evidence by the loss in fluorescence. When challenged with the RPA+Rtt105 complex, no FE is observed. Thus, Rtt105 blocks FE activity of RPA and thus likely contributes to the formation of stable RPA nucleoprotein filaments.

**Figure 6.**
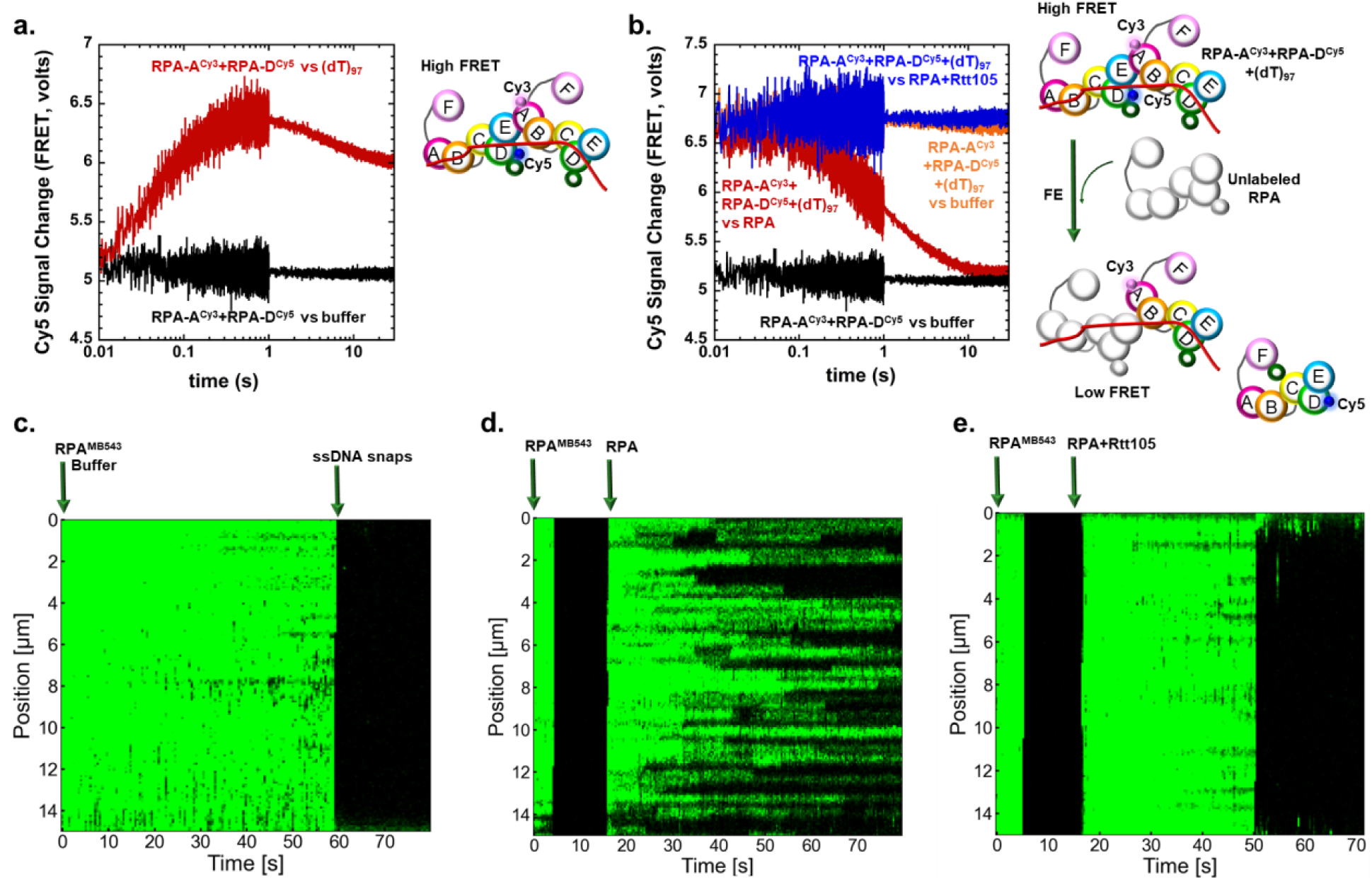
Rtt105 inhibits facilitated exchange activity of RPA. **a)** A high FRET complex is formed when multiple molecules of RPA-DBD-A^Cy5^ and RPA-DBD-D^Cy3^ bind to a (dT)_97_ substrate (red). FRET-induced Cy5 fluorescence is not observed in the absence of DNA (black). **b)** Facilitated exchange (FE) activity was measured by mixing preformed [RPA-DBD-A^Cy5^:RPA-DBD-D^cy3^:(dT)_97_] complexes with unlabeled RPA (red) resulting in a loss of Cy5 fluorescence. Rtt105 inhibits the FE activity as preformed Rtt105:RPA complexes do not perturb the Cy5 signal (blue). Control experiments in the absence of unlabeled RPA (orange) or DNA (black) are also shown as reference for the high and low FRET-induced Cy5 fluorescence states. Models for FE are denoted on the right. **c)** C-trap experiments to investigate FE of RPA were performed on ssDNA that was generated by mechanically unfolding lambda dsDNA (∼48.5kbp) in the optical trap and subsequently incubated with RPA-DBD-D^MB543^ (10 nM) for 15-20s and stably formed fluorescent RPA coated ssDNA nucleoprotein filaments are observed. **d)** The fluorescent RPA nucleoprotein filament was subsequently moved to a channel containing unlabeled RPA (10 nM) and after a 10s incubation in the dark FE can be observed. Spots where loss of fluorescence is observed are events where the unlabeled RPA have replaced fluorescent RPA molecules. **e)** When challenged with unlabeled RPA:Rtt105, no loss in fluorescence is observed. Thus, Rtt105 inhibits the FE activity of RPA.

### Complex formation with Rtt105 negatively regulates interactions with RPA-interacting proteins like Rad52

Since Rtt105 configurationally staples RPA and restricts the DBDs and PIDs we wondered if the protein-protein interaction activities of RPA were regulated by Rtt105. This could prevent spurious RPA interactions with its myriad binding partners upon nuclear import from the cytoplasm and add to the functions of Rtt105 in genomic integrity. To test this hypothesis, we tested whether Rtt105 influenced RPA binding to Rad52, a mediator protein that physically interacts with RPA and functions to promote homologous recombination (HR) (Park et al., 1996). SEC has limitations in resolving larger molecular weight complexes, thus we used analytical ultracentrifugation (AUC) to monitor complex formation. Rtt105, RPA, Rad52 all sediment as distinct complexes (**Figure 7a**). RPA and Rad52 form a complex in the absence or presence of ssDNA (dT)_35_ (**Figure 7a**). We and others have previously shown that Rad52 interacts with RPA on and off the DNA and these data agree with our knowledge of these interactions (Park et al., 1996; Plate et al., 2008; Pokhrel et al., 2019). The only key difference is the oligomeric state of yeast Rad52 which is widely considered to be a heptamer (Shinohara et al., 1998). In our ongoing cryoEM studies, *S. cerevisiae* Rad52 is a homodecamer (Deveryshetty and Antony, unpublished data) and the stoichiometries denoted here reflect this finding. In the absence of ssDNA, Rtt105 remains as a complex with RPA and prevents interaction with Rad52 (**Figure 7b**). Thus, Rtt105 functions as a negative regulator of RPA-Rad52 interactions in the absence of DNA. In the presence of ssDNA Rtt105 is released and formation of RPA-DNA, RPA-Rad52, and RPA-Rad52-DNA complexes are observed (**Figure 7c**). These data suggest that ssDNA acts as a licensing agent to release Rtt105 from RPA and promote RPA interactions with Rad52, and likely with other RPA-interacting proteins (**Figure 7d**).

**Figure 7:**
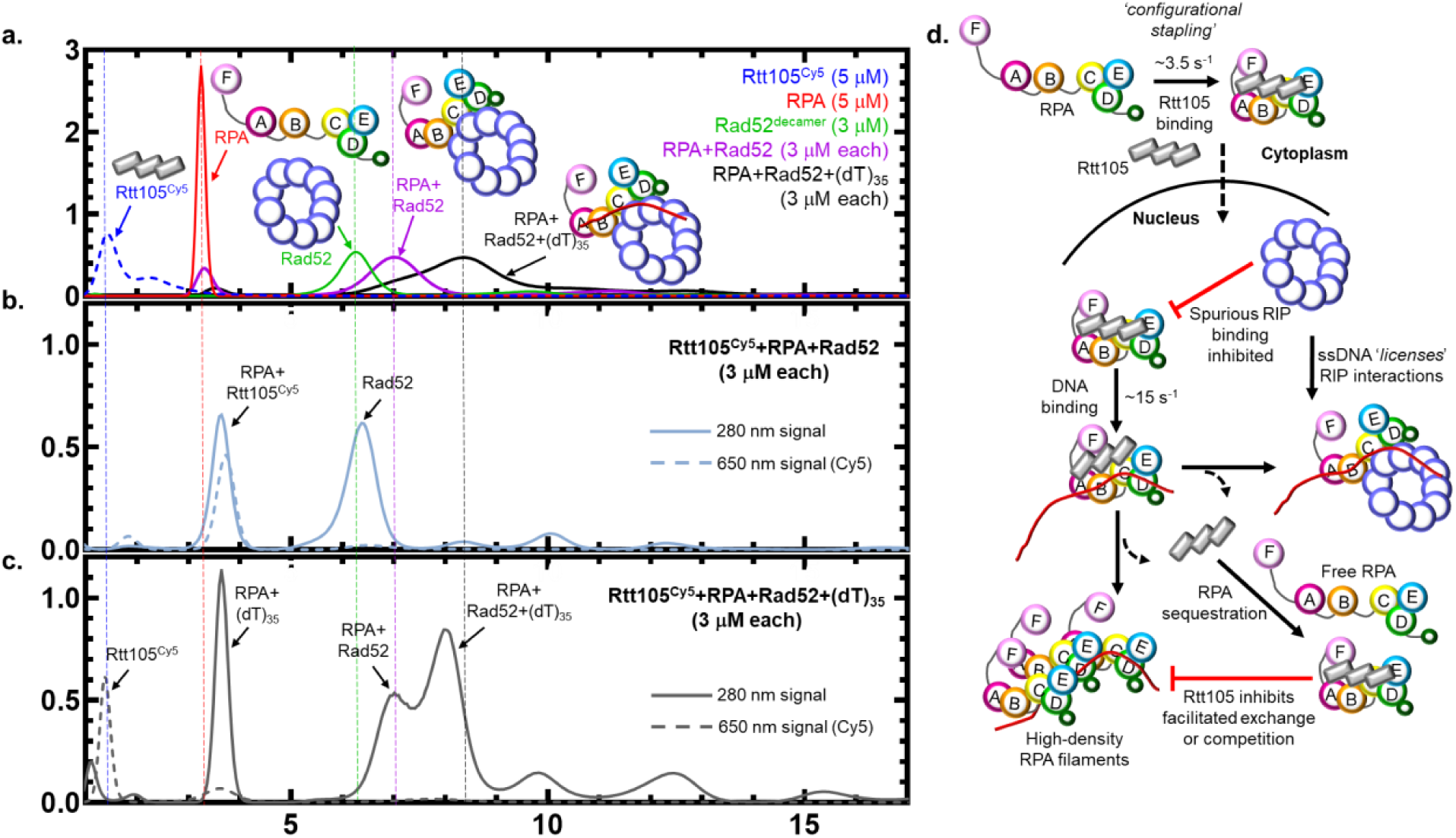
Rtt105 inhibits spurious RPA-Rad52 interactions in the absence of ssDNA. **a)** Analytical ultracentrifugation (AUC) analysis of RPA, Rtt105^Cy5^, RPA, and the RPA-Rad52 and RPA-Rad52-(dT)_35_ complexes show them sedimenting as distinct complexes. The 280 nm signals were monitored for all the proteins and complexes except Rtt105^Cy5^ which was monitored at 650 nm (dashed line). The horizontal dotted lines serve as a visual guide for the sedimentation positions of the denoted proteins across all the AUC experiment panels. **b)** Rtt105^Cy5^, RPA, and Rad52 were mixed in equimolar concentrations and analyzed. In the presence of Rtt105, Rad52 does not interact with RPA and sediments independently while RPA and Rtt105 remain as a complex. **c)** When an equimolar amount of ssDNA [(dT)_35_] is present, Rtt105 is displaced and sediments similar to free Rtt105. RPA and Rad52 partition into multiple RPA+DNA, RPA+Rad52, and RPA+Rad52+DNA complexes. Importantly, Rtt105 is not part of most of these complexes. A small fraction of the Rtt105 is present along with the RPA-DNA complex. **d)** Rtt105 configurationally staples RPA through interactions with multiple domains including the DNA binding and protein interaction domains. The complex is shuttled into the nucleus where promiscuous interactions with RPA binding proteins such as Rad52 are inhibited by Rtt105 in the absence of ssDNA. Availability of ssDNA triggers configurational and conformational changes within the RPA-Rtt105 complex and this higher order complex can interact with RIPs such as Rad52 or drive assembly of multiple RPA-bound DNA structures. Both these scenarios drive dissociation of Rtt105. Rtt105 can also sequester free RPA to prevent untimely facilitated exchange (FE) events as the RPA-Rtt105 complex cannot perform FE.

## Discussion

Rtt105 functions akin to a chaperone-like protein to transport RPA into the nucleus, and deletion of Rtt105 reduces RPA availability for DNA metabolic roles including DNA repair and recombination (Corda et al., 2021; Li et al., 2019; Li et al., 2018; Wang et al., 2021). Previous studies have proposed a model where Rtt105 stretches the many domains of RPA to enhance DNA binding (Li et al., 2019). Here, we present a contrasting model for Rtt105 mediated regulation of RPA. First, our findings show extensive contacts between Rtt105 and RPA that promotes a compaction of the four DNA binding and two protein-interaction domains of RPA. The CL-MS and HDX MS analysis show configurational and conformational changes in all domains and a few of the intervening flexible linkers. While a structure of Rtt105 or the RPA-Rtt105 complex is not yet available, the AlphaFold prediction renders a helical protein that could be stretched over a large radius (**Figure 1b**). Thus, we propose that configurational stapling compacts RPA through extensive contacts with Rtt105 and promotes nuclear import (**Figure 7d**). Rtt105 is an extensively charged protein and stretches of negatively charged patches are observed and likely binds close to the positively charged DNA binding pockets in RPA. In support of this model, we see differences in the DNA binding properties of the Rtt105-RPA complex.

While Rtt105 does not bind directly to DNA, Rtt105 complexed with RPA influences the kinetics and conformations of the DBDs on ssDNA. Positioning of site-specific fluorophores on either DBD-A or DBD-D enables us to quantify how each of these domains bind to DNA and are remodeled. Rtt105 reduces the rate of ssDNA binding by DBD-D by ∼2-fold but does not change the final configuration of this domain on DNA. In contrast, Rtt105 does not change the rate of ssDNA binding to DBD-A, but significantly alters the final configuration. Thus, we envision Rtt105 situated on both the Tri-C and FAB halves of RPA in the absence of ssDNA as supported by the XL-MS analysis (**Figure 4**). The region close to Tri-C occludes the DNA binding site in DBD-D (possibly in DBD-C as well) and is remodeled in the presence of DNA. The Rtt105 region close to the FAB region likely does not block DNA binding but restrains the configurational freedom of those domains. In the [Rtt105-RPA-(dT)_35_] complex, ssDNA must be bound close to the Tri-C region. Further evidence for this model arises from the dissociation of Rtt105 on ssDNA longer than (dT)_35_ where multiple RPA can bind. Here, DBD-A (RPA70) from one RPA contacts the OB-E (RPA14) of the neighboring RPA (Yates et al., 2018).This interaction likely triggers the dissociation of Rtt105. Moreover, Rtt105 also promotes formation of RPA nucleoprotein filaments that have a higher density of RPA molecules bound. This property renders the RPA nucleoprotein filament more stable and explains the additional stretching of ssDNA observed in DNA curtain experiments (Li et al., 2018). In the cell, formation of such RPA nucleoprotein platforms triggers the DNA damage response (Marechal and Zou, 2015) and Rtt105 might contribute to such functions by not releasing RPA until longer stretches of ssDNA become exposed.

Contradictory to earlier studies, we do not see a difference in the K_D_ for RPA ssDNA interactions in the presence of Rtt105. This is not surprising as RPA will be macroscopically bound with high affinity to DNA while select DBDs can undergo dynamic rearrangements (Caldwell and Spies, 2020). Moreover, because of the high ssDNA binding affinity (K_d_<10^−10^M), enhancements of DNA binding affinity cannot be measured using traditional bulk ensemble measurements. In both EMSA and anisotropy measurements we do not see an Rtt105 induced enhancement of RPA-ssDNA interactions. Our *in vivo* studies show a distinction between functions associated with Rtt105 and DNA interactions of RPA. Loss of Rtt105 (*Δrtt105*) or reduction in RPA DNA interactions (*rfa1 zm1, zm2*, or *t33* mutations) shows defects in cell growth and a combination of *Δrtt105* and *rfa1* mutations result in severe growth defects. The additive nature of these phenotypes again suggests that Rtt105 does not directly influence the DNA binding properties of RPA, but regulates it through either affecting nuclear cytoplasmic shuttling, RIP interactions, or processes such as facilitated exchange. Rtt105 contributes to the levels of RPA in the nucleus, however, no severe defects in global DNA synthesis are observed under non-stressed conditions (Corda et al., 2021). In contrast, at perturbed replication forks, the lack of Rtt105 becomes important for RPA loading (Corda et al., 2021). In this scenario, longer ssDNA intermediates are generated and thus, the inhibition of FE by Rtt105 could explain the formation of more stable RPA filaments.

Since RPA interactions with Rtt105 occur in the cell well before the availability of ssDNA, we hypothesize that posttranslational modifications of RPA (and/or Rtt105), such as phosphorylation by kinases, might influence the interaction. In support of this idea, a phosphomimetic of RPA carrying a S->D substitution at position 178 in RPA70 (RPA^S178D^) shows reduced binding to Rtt105 (**Supplemental Figure S16**). Rtt105 has also been implicated in the resolution of G-quadruplex structures by RPA (Maestroni et al., 2020). *In vitro*, we do not see any direct Rtt105 enhancement of RPA promoted G-quadruplex unwinding (**Supplemental Figure S17**). Thus, we propose that Rtt105 regulates the activity of RPA by controlling its nuclear transport, inhibiting RIP interactions in the absence of ssDNA, and regulating the availability of free RPA.

Finally, in higher eukaryotes, RPAIN (RPA interacting protein) or RIPα (RPA interacting protein α) is proposed to be the functional ortholog of Rtt105 as they share no similarity in sequence (Wang et al., 2021). The AlphaFold predicted structure of RPAIN shows a reasonable degree of resemblance to Rtt105 (**Supplemental Figure S18**). RPAIN has also been proposed to function by enhancing the DNA binding activity of human RPA (hRPA) (Wang et al., 2021). Similar to our observations for Rtt105 and yeast RPA, we see complex formation between hRPA and RPAIN, but do not observe a stimulation of DNA binding properties (**Supplemental Figure S18**). Interestingly, shorter DNA (dT)_35_ can promote the dissociation of the RPAIN-hRPA complex, and future work will focus on deciphering the differences in the mechanism of action of RPAIN.

## Materials and Methods

### Reagents and Buffers

Chemicals were purchased from Sigma-Millipore Inc., Research Products International Inc. and Gold Biotechnology Inc. Fluorescent and unlabeled oligonucleotides were synthesized by Integrated DNA Technologies. Enzymes for molecular biology were purchased from New England Biolabs. Resins for protein purification were sourced from GE-Cytiva Life Sciences Inc. Fluorophores for protein labeling were from Click Chemistry Tools Inc.

### Plasmids for protein overproduction

An RSF-Duet1 plasmid coding for Rtt105 with a N-terminal 6x-polyhistidine tag was used. Mutations in *Rtt105* were introduced using site-directed mutagenesis. Plasmids for RPA and 4-azidophenylalanine (4AZP) incorporation were as described (Kuppa et al., 2021; Pokhrel et al., 2019; Pokhrel et al., 2017).

### Purification of Rtt105 and RPA

Rtt105 was overproduced in Rosstta-2 PlysS *E. coli* cells and purified as described (Li et al., 2018). *Saccharomyces cerevisiae* RPA was purified as described (Pokhrel et al., 2017). Non-canonical amino acid (4AZP) incorporation based fluorescently labeled RPA were generated as described (Kuppa et al., 2021; Pokhrel et al., 2019; Pokhrel et al., 2017). Concentration of Rtt105 was determined spectroscopically using ε_280_ =18,450 M^-1^cm^-1^. Rtt105 was flash frozen and stored at -70 °C. Concentration of unlabeled and labeled RPA was measured spectroscopically using ε_280_ =98,500 M^-1^cm^-1^. Labeling efficiency for fluorescent RPA was calculated as described using absorption values measured at 280 nm and ε_280_= 98500 M^-1^cm^-1^for RPA, at 550 nm with ε_550_= 105,000 M^-1^cm^-1^for RPA-MB543, at 555 nm with ε_555_= 150,000 M^-1^cm^-1^for RPA-Cy3, and at 650 nm with ε_650_= 250,000 M^-1^cm^-1^ for RPA-Cy5 versions (Kuppa et al., 2021; Pokhrel et al., 2019).

### Generation of fluorescently-labeled Rtt105

Rtt105 has two consecutive Cys residues at positions 12 and 13, respectively. Either or both Cys residues can be substituted with Ser without loss of binding to RPA (**Supplemental Figure S2**). We used Cys-12 for attachment of fluorophores using maleimide chemistry and converted Cys-13 to Ser. This version of Rtt105 (Rtt105^C13S^) was purified similar to the wild-type protein and ∼5 ml of 100 μM Rtt105^C13S^ was dialyzed extensively in labeling buffer (30 mM HEPES, pH 7.8, 200 mM KCl, 0.25 mM EDTA, pH 8.0, 0.01 % Tween-20, and 10 % v/v glycerol) to remove TCEP-HCl from the storage buffer. After 4 buffer exchanges, a 1.5-fold molar excess of Cy5-malemide dye was added to the dialyzed protein and incubated at 4 °C for 3 hours. The reaction was then quenched with 0.5 % β-mecraptoethanol (βME). Excess dye was separated from the protein using a Biogel-P4 column (Bio-rad laboratories). The concentration of Rtt105 calculated spectroscopically using extinction coefficient ε_280=_ 18,450 M^-1^cm^-1^ and labeling efficiency was calculated using the Cy5 absorbance signal and ε_650_= 250,000 M^-1^cm^-1^. Rtt105 absorbance values at 280 nm were also corrected for minor signal interference from Cy5 by measuring the percent contribution of free Cy5.

### Yeast strains and genetic techniques

Standard procedures were used for cell growth and media preparation. Strains used are provided in the table below and are isogenic to W1588-4C, a *RAD5* derivative of W303 *(MATa ade2-1 can1-100 ura3-1 his3-11,15, leu2-3, 112 trp1-1 rad5-535)* (Zhao and Blobel, 2005). Gene deletion to generate *rtt105*Δ strain was performed following standard PCR based method. Standard yeast genetic procedures were used for tetrad analyses and at least two biological duplicates were used for each genotype.

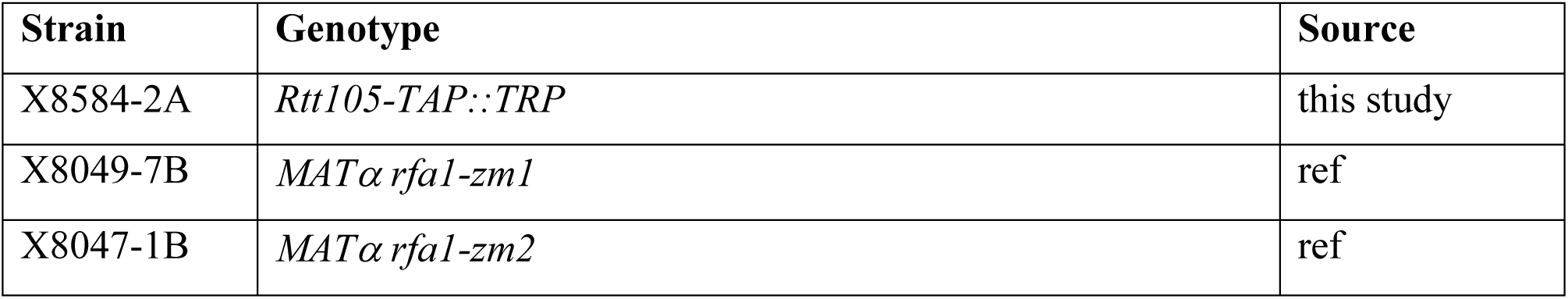
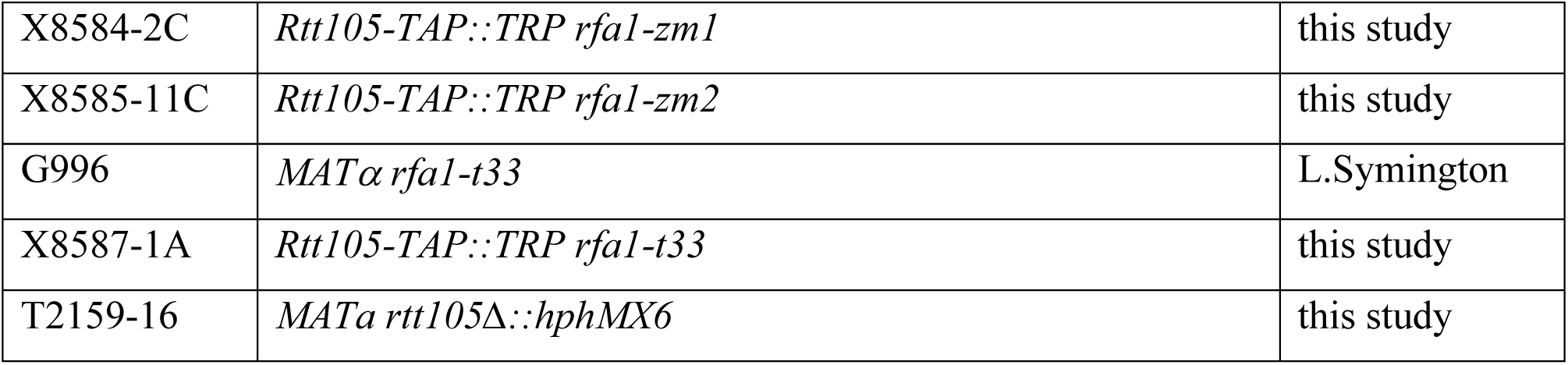

### Co-Immunoprecipitation

Yeast cells from log phase cultures growing in YPD were harvested and lysed by bead beating in TMG-140 buffer (10 mM Tris-HCl, pH 8.0, 4 mM MgCl_2_, 10 % v/v glycerol, 140 mM NaCl, 0.1 mM EDTA, 0.5 % Tween20, 1 mM DTT and Roche cOmplete-Ultra EDTA free protease inhibitor). DNA was digested by incubation with benzonase for 30 minutes at 4 °C. Lysates were cleared by centrifugation and incubated in TMG-140 buffer with IgG sepharose beads for 2 hours at 4 °C. After incubation, beads were washed with TMG-140 and proteins were eluted with Laemmli buffer. Proteins were separated on gradient gels followed by western blotting with antibodies against Rfa1 (a kind gift from Dr. Steven Brill at Rutgers University).

### Secondary structure analysis using circular dichroism (CD)

CD measurements were used to compare the secondary structures of Rtt105 and the Cys mutant variants of Rtt105. A nitrogen-fused observation chamber with a cell pathlength of 1 mm was used. All CD traces were obtained between 200-260 nm at 20 °C on a Chirascan CD spectrometer (Applied Photophysics). 600 nM of Rtt105^WT^, Rtt105^C13S^, Rtt105^C12S^, Rtt105^CCSS^ in CD reaction buffer (5 mM Tris-Cl pH 7.8, 100 mM KCl, 5 mM MgCl_2_, and 6% v/v glycerol) was used to obtain CD spectra. The results were collected using 1 nm step size, 1 nm bandwidth and 5 traces were averaged.

### Analysis of complex formation using size exclusion chromatography (SEC)

600 μl of the noted concentrations of RPA, Rtt105, or the complex in the absence or presence of equimolar amounts of DNA were resolved on a 10/300 Superose 6 Increase column using an AKTA-pure FPLC system. Protein and protein-DNA complexes were incubated at 4 °C for 10 min before analysis. Resolution was performed using Rtt105-SEC buffer (30 mM HEPES, pH 7.8, 100 mM KCl, 1 mM TCEP-HCl, and 10% v/v glycerol). A total of 30 ml elution volume was collected as 0.5 ml fractions and further analyzed on 10% SDS-PAGE.

### Measurement of RPA-ssDNA interactions using electrophoretic mobility band shift analysis (EMSA)

10 nM 5′-Cy5-(dT)_30_ ssDNA was incubated with indicated amounts of RPA at 25 °C in EMSA buffer (25 mM Tris-Cl, pH 7.5, 200 mM NaCl, 5 mM MgCl_2_, 5 % v/v glycerol, and 0.05 % Tween-20) for 10 min. For experiments performed in the presence of Rtt105, RPA and Rtt105 were premixed in equimolar ratios at 25 °C for 5 min before DNA was introduced. The reaction mixture (20 μl) was mixed with 10 μl of 70 % v/v glycerol, mixed, and loaded onto a 6 -15 % bisacrylamide gradient gel and resolved using 1x TBE buffer. The gels were scanned using an iBright 1500 imager (Thermo Fischer Inc.) and the Cy5 fluorescence associated with ssDNA (unbound fraction) or ssDNA+protein (bound fraction) bands was background subtracted and quantified with the associated iBright software. The mean values and standard deviation from three independent experiments were plotted for analysis. K_D_ was estimated by nonlinear least squares fitting to the following equation (Eq. 1):

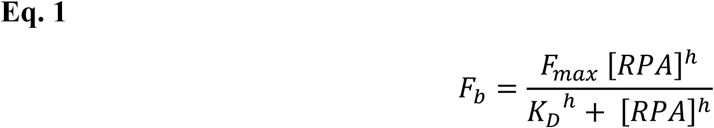

*F*_*b*_ is the fraction of bound ssDNA determined from fluorescence intensities of two bands, namely:

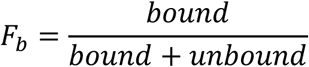

*F*_*max*_ is the fraction of bound ssDNA at saturating protein concentration, *K*_*D*_ is the apparent dissociation constant, [*RPA*] is concentration of RPA or RPA+Rtt105 (1:1) in each well, and *h* is the Hill coefficient.

*Measurement of RPA and ssDNA binding in presence or absence of Rtt105 using fluorescence anisotropy*. 5’-FAM-(dT)_35_ ssDNA was diluted to 10 nM in 1x RPA reaction buffer (30 mM HEPES pH 7.8, 100 mM KCl, 6 % v/v glycerol, 5 mM MgCl_2_, and 1 mM βME). 180 μl of this working stock was added to a 3 mm pathlength quartz cuvette (Starna Cells Inc.) and the temperature was maintained at 23 °C. Fluorescence anisotropy of the FAM-labeled ssDNA was measured using a PC1 spectrofluorometer (ISS Inc.). Samples were excited at 488 nm and the resulting emission was collected using a 520 nm bandpass emission filter. Five anisotropy readings were acquired and averaged before or after stepwise addition of RPA alone, or 1:1 stock of RPA and Rtt105 (each protein at 5 μM). Four independent experiments were performed, and the mean and SEM were estimated from G-factor corrected values. K_D_ was determined after fitting baseline subtracted anisotropy values with a model for one site specific binding with Hill slope as defined by Eq.2 (below) using GraphPad Prism 9.

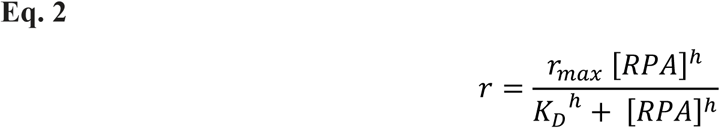

Where, *r* is the measured G-factor corrected anisotropy value, *r*_*max*_ is the maximum anisotropy when 100% of ssDNA is complexed with RPA, *K*_*D*_ is the apparent dissociation constant, [*RPA*] is the concentration RPA in the cuvette after each successive addition, and *h* is the Hill coefficient.

For the anisotropy experiments where the binding density of RPA was measured, 30 nM 5′-FAM-(dT)_35_ or 5′-FAM-(dT)_70_ were taken in 1x RPA reaction buffer in a 10 mm pathlength quartz cuvette (Firefly Sci) with stirring. RPA alone or RPA + Rtt105 (1:1) were titrated and after an incubation period of 3 min fluorescence anisotropy was measured and plotted against the concentration of proteins. RPA and Rtt105 were mixed at 1:1 molar ratio (1.1 μM each) in 1x RPA reaction buffer and incubated on ice for 30 min prior to each experiment. The experiments were repeated three times and the mean and SEM were estimated from G-factor corrected values. The saturation points were taken as the intersection of biphasic or triphasic curves from the linear fits of the initial data points reflecting the change in anisotropy upon binding of sub-saturating amounts of proteins. For (dT)_35_, the first dotted line represents a stoichiometry of 1:1, and the other represents 2.7:1. For (dT)_70_, first dotted line represents a stoichiometry of 2:1, and the other represents 4.7:1. However, note that stoichiometry estimates with (dT)_70_ is probably an underestimate because an anisotropy value of 0.19 is close to the limiting value of anisotropy measurements for fluorescein.

### Crosslinking Mass Spectrometry (XL-MS) analysis

Stock solutions of Rtt105 (13.4 mg/mL) and RPA (1.77 mg/mL) were diluted to 0.89 mg/mL and 0.3752 mg/mL, respectively in buffer (30 mM HEPES, 200 mM KCl, pH 7.8 and incubated together for 30 minutes. The diluted proteins were reacted with 5 mM bis(sulphosuccinimidyl)suberate (BS3) and 20 µL of the sample was taken at various time points (0, 15 and 30 minutes) and immediately quenched with 2 µL of 1M ammonium acetate. Quenched samples were diluted with 1.5X Laemmli gel loading buffer to a final volume of 40 µL, vortexed, and heated to 100 °C for 5 min and resolved on 4-20% (w/v) gradient SDS-PAGE gels (Bio-Rad) with Tris-glycine buffer. Gels were stained with Gelcode blue safe protein stain (Thermo Scientific). Gel bands were excised for protein identification and analysis. Excised bands were destained with a 50 mM ammonium bicarbonate and 50% acetonitrile mixture and reduced with a mixture of 100 mM DTT and 25 mM ammonium bicarbonate for 30 minutes at 56 °C. The reaction was subsequently exchanged for the alkylation step with 55 mM iodoacetamide and 25 mM ammonium bicarbonate and incubated in the dark at room temperature for 25 min. The solution was then washed with the 50 mM ammonium bicarbonate and 50 % acetonitrile mixture. The gel pieces were then first dehydrated with 100 % acetonitrile and then rehydrated with sequence grade trypsin solution (0.6 µg, Promega) and incubated overnight at 37 °C. The reaction was quenched with 10 µL of 50% acetonitrile and 0.1% formic acid (FA, Sigma) and transferred to new microfuge tubes, vortexed for 5 min, and centrifuged at 15,000 rpm for 30 min. Samples were transferred to mass spectrometry vials and quantitated by LC-MS as described for peptide identification (Burns et al., 2017; Kim et al., 2022). Peptides were identified as previously described (Berry et al., 2018) using MassHunter Qualitative Analysis, version 6.0 (Agilent Technologies), Peptide Analysis Worksheet (ProteoMetrics LLC), and PeptideShaker, version 1.16.42, paired with SearchGUI, version 3.3.16 (CompOmics). Crosslinks were then determined using Spectrum Identification Machine (SIMXL 1.5.5.2).

### Hydrogen-Deuterium Exchange mass spectrometry (HDX-MS) analysis

Stock solutions of RPA (13.4 mg/mL) and Rtt105(1.77 mg/mL) were mixed in the presence or absence of (dT)_35_ ssDNA in a 1:1.2 ratio. Reactions were diluted 1:10 into deuterated reaction buffer (30 mM HEPES, 200 mM KCl, pH 7.8). Control samples were diluted into a non-deuterated reaction buffer. At each time point (0, 0.008, 0.05, 0.5, 3, 30 h), 10 µL of the reaction was removed and quenched by adding 60 µL of 0.75 % formic acid (FA, Sigma) and 0.25 mg/mL porcine pepsin (Sigma) at pH 2.5 on ice. Each sample was digested for 2 min with vortexing every 30 s and flash-frozen in liquid nitrogen. Samples were stored in liquid nitrogen until the LC-MS analysis. LC-MS analysis of RPA was completed as described (Patterson et al., 2020). Briefly, the LC-MS analysis of RPA was completed on a 1290 UPLC series chromatography stack (Agilent Technologies) coupled with a 6538 UHD Accurate-Mass QTOF LC/MS mass spectrometer (Agilent Technologies). Peptides were separated on a reverse phase column (Phenomenex Onyx Monolithic C18 column, 100 × 2 mm) at 1 °C using a flow rate of 500 μl/min under the following conditions: 1.0 min, 5% B; 1.0 to 9.0 min, 5 to 45% B; 9.0 to 11.8 min, 45 to 95% B; 11.8 to 12.0 min, 5% B; solvent A = 0.1 % FA (Sigma) in water (Thermo Fisher) and solvent B = 0.1% FA in acetonitrile (Thermo Fisher). Data were acquired at 2 Hz s^-1^ over the scan range 50 to 1700 m/z in the positive mode. Electrospray settings were as follows: the nebulizer set to 3.7 bar, drying gas at 8.0 L/min, drying temperature at 350 °C, and capillary voltage at 3.5 kV. Peptides were identified as previously described (Berry et al., 2018) using MassHunter Qualitative Analysis, version 6.0 (Agilent Technologies), Peptide Analysis Worksheet (ProteoMetrics LLC), and PeptideShaker, version 1.16.42, paired with SearchGUI, version 3.3.16 (CompOmics). Deuterium uptake was determined and manually confirmed using HDExaminer, version 2.5.1 (Sierra Analytics). Heat maps were created using MSTools (Kavan, 2010).

### MB543 fluorescence quenching assay to estimate binding affinity between RPA and Rtt105

RPA-DBD-A^MB543^, RPA-DBD-D^MB543^, or F-A-B-DBD-A^MB543^ were diluted to 200 nM in 1x RPA reaction buffer. 180 μl of either fluorescent protein was added to a 3 mm path length quartz cuvette (Starna Cells Inc.) and maintained at 23 °C in a PC1 spectrofluorometer (ISS Inc.). The MB543 dye, conjugated to either DBD-A or DBD-D domain of RPA, was excited at 535 nm and the resulting fluorescence emission spectra were collected between 558 nm to 578 nm with the λ_max_ at 568 nm. Unlabeled Rtt105, diluted to a 4 μM stock in RPA reaction buffer, was added to the cuvette in a stepwise manner, mixed, and incubated for 3 minutes to achieve equilibrium before emission spectra were measured. Emission scans were recorded twice from at least two cuvettes, and the experiment was repeated three to four times. The fluorescence at λ_max_ from 3-4 trials was corrected for stepwise dilution of sample (<8%), normalized to the initial fluorescence (fluorescence intensity in the absence of Rtt105), and plotted as mean and SEM. The fluorescence intensity values were transformed to fraction quenched versus Rtt105 concentration and fitted to a quadratic equation (Eq. 3, below) using non-linear least squares regression, accounting for ligand depletion, to yield an apparent equilibrium constant (K_D_). The dilution factor corrected RPA-DBD-D^MB543^ fluorescence remained nearly constant (within error) and served as a control for change in fluorescence due to photobleaching alone.

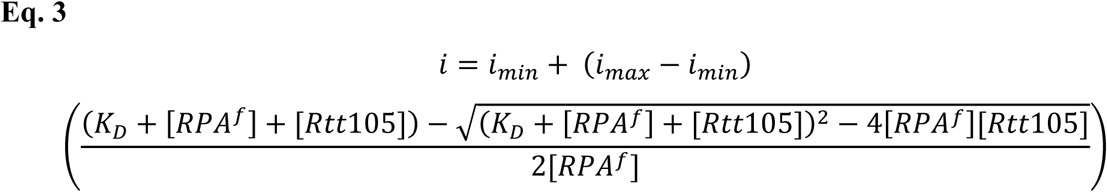

Where, *i*, is the measured fluorescence intensity, *i*_*min*_, & *i*_*max*_ are minimum and maximum values of the fluorescence intensity of 100% free, and 100% bound RPA determined from the fit, respectively. [*RPA*^*f*^], is the concentration of fluorescent RPA taken in the cuvette and the value is constrained at 200 nM. However, note that due to the stepwise addition of Rtt105 there is a 5% dilution by the end of the measurement. [*Rtt*105] is the dilution factor corrected concentration of Rtt105 in the cuvette in nM, and *K*_*D*_ is the dissociation constant determined from the fit.

### Measurement of DNA binding kinetics using stopped flow fluorescence

All stopped flow experiments were performed on a SX20 instrument (Applied Photophysics Inc.) at 25 °C in 1x RPA reaction buffer. Protein and or DNA reactions from individual syringes were rapidly mixed and fluorescence data were collected. The respective mixing schemes are denoted by cartoon schematics within the figure panels. Seven to eight individual shots were averaged for each experiment. All experiments were repeated a minimum of 3 times and SEM from the individual fits are noted in the figure legends. For the FRET experiments, samples were excited at 535 nM (Cy3 wavelength) and Cy5 emission was captured using a 645 nm long-pass filter. For the RPA-Rtt105 interactions, RPA-DNA and Rtt105-RPA-DNA interactions, experiments were performed with 100 nM each of RPA, Rtt105, and (dT)_35_ or (dT)_40_ ssDNA substrates (1:1:1 ratio). For facilitated exchange stopped flow experiments, 200 nM RPA-DBD-A^Cy5^ and 200 nM RPA-DBD-D^Cy3^ were premixed with 120 nM (dT)_97_ and shot against unlabeled RPA (500 nM) or the RPA-Rtt105 complex (500 nM each).

### Forster resonance energy transfer (FRET) measurement of protein-DNA and protein-protein complexes

RPA-DBD-D^Cy3^ and Rtt105^Cy5^ were mixed in 1x RPA reaction buffer at 1:1 ratio such that the final concentration of each protein was 200 nM. The complex was incubated on ice for 30 minutes and then transferred to a 3 mm pathlength cuvette maintained at 23 °C in the PC1 spectrofluorometer. The sample was excited at 535 nM and the resulting fluorescence between 550 nm to 700 nm was collected as a FRET spectrum. Next, ssDNA of different lengths (dT_x_); where x = (dT)_8_, (dT)_15_, (dT)_25_, (dT)_35_, (dT)_45_, (dT)_54_, (dT)_64_, (dT)_70_, or (dT)_84_ were added in a stepwise manner to the cuvette, mixed, and incubated for 3 minutes at 23 °C before FRET spectra was again recorded after each addition. Raw spectra were corrected by incorporating the estimated dilution factor and then area normalized to account for any fluctuations in lamp intensity. Finally, ratiometric FRET was calculated as defined by Eq. 4:

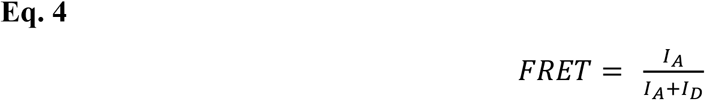

Where, I_A_ and I_D_ are acceptor and donor fluorescence emission intensities at respective λ_max_ = 573, and 673 nM, respectively.

### C-Trap optical tweezer analysis of RPA-Rtt105 interactions

48.5 kbp Lambda DNA construct was prepared with three biotins on either end of DNA. Lambda DNA was purchased from Roche Inc. Short oligos to anneal to the sticky ends were purchased from Integrated DNA Technologies Inc. DNA were stored in TE buffer (10 mM Tris-HCl, pH 8.0 and 0.1 mM EDTA). Optical trap experiments were performed using a commercial dual optical trap combined with confocal microscopy and microfluidics [C-trap] from Lumicks BV Inc. Streptavidin coated polystyrene particle beads of average size 4.8 µM [0.5 % w/v] (Spherotech Inc.) were diluted 1:250 in 1X PBS and 1-2 nM of DNA were made in 1X PBS. DNA was captured between two streptavidin beads and mechanically denatured by moving one bead to the overstretched region to create ssDNA. ssDNA was confirmed by fitting force-distance (FD) curve to Freely Jointed Chain model [FJC] (contour length 48.5 kbp / 16.49 um; persistence length 46 nm; stretch modulus 1000 pN) in real time. DNA was held for 5 s in the fully ssDNA state then returned to a 5 pN tension on ssDNA position for the fluorescence experiments. RPA-DBD-D^MB543^ in storage buffer (30 mM HEPES pH 7.8, 200 mM KCl, 0.02 % Tween-20, 10 % glycerol, and 0.2 mM EDTA) was added to the DNA in the absence or presence of Rtt105^Cy5.^ Rtt105^Cy5^ was kept in storage buffer with 1 mm TCEP-HCl. Both proteins were diluted to 1 nM with experimental buffer (30 mM HEPES pH 7.8, 100 mM KCl, 6 % Glycerol, 5 mM MgCl_2_ and incubated together (for the experiments where Rtt105-RPA complexes were tested) at 1:1 molar ratio (10 pM final concentration each). Imaging buffer 0.8 % (w/v) dextrose, 165 U/mL glucose oxidase, 2170 U/mL catalase, and 2-3 mM Trolox was used to increase the fluorescence lifetime of the fluorophores. Imaging settings were 2-3 ms exposure time (per pixel), red excitation 638 nm, and green excitation 561 nm. Data was analyzed using custom python script [Pylake API from Lumicks] and code is available upon request.

### Analytic Ultracentrifugation (AUC) analysis

AUC sedimentation velocity experiments were performed on an Optima analytical ultracentrifuge (Beckman-Coulter Inc.) using an An-50Ti rotor at 40,000 rpm at 20 °C. Proteins and DNA either alone or in complex were dialyzed against 30 mM HEPES, pH 7.8, 100 mM KCl, 10 % glycerol, and 1 mM TCEP before each experiment. Concentrations used for the experiments are mentioned in the appropriate figures. Sample (380 μL) and buffer (400 μL) were filled in each chamber of a 2-sector charcoal quartz cell. Absorbance was monitored at 280 nm and/or 650 nm. Since the absorbance signal from Rtt105 was low Rtt105^Cy5^ was used and tracked at 650 nm. Scans were recorded at 3 min intervals. The density and viscosity of the buffer at 20 °C was calculated using SEDNTERP. Continuous distribution (c(s)) model was used to fit the data in SEDFIT (Schuck, 2000).

## Acknowledgements

The authors thank members of our respective research laboratories for critical reading of the manuscript.

## Author Contributions

SK, JD, RC, NP, VK and MKS purified proteins, generated fluorescent protein, designed and performed the steady-state and pre-steady state experiments. JM, AP, SK and BB performed and analyzed the HDX-MS and XLMS experiments. SP and TH performed and analyzed the C-trap experiments. ND and XZ performed and analyzed the *in vivo* experiments. S.S. and H.B. performed and analyzed single molecule FRET measurements of GQ-unwinding. EA directed the projected, designed experiments, and composed the manuscript. All authors contributed to manuscript preparation.

## Data availability

All data are contained within the manuscript. Plasmids used for protein overexpression are available upon request. Code for C-trap data analysis is available upon request.

## Funding and additional information

This work was supported grants from the National Institutes of Health (R01 GM130746 and R01 GM133967) to E.A., (R15 GM123443) to H.B., (R35 GM122569) to T.J. and (R01 GM131058) to X.Z. Funding for Proteomics, Metabolomics and Mass Spectrometry Facility at MSU was made possible in part by the MJ Murdock Charitable Trust and NIGMS of the National Institutes of Health under Award Number P20 GM103474. The analytical ultracentrifuge experiments were supported by an instrumentation grant from the Office of the Director, National Institutes of Health (S10 OD030343) to E.A. M.K.S. was supported by a summer undergraduate research supplement from the National Institutes of Health (R01 GM130746-04S1 to E.A.). Circular dichroism and fluorescence experiments were performed on instruments funded through supplement grants from NIG-NIGMS (R01 GM133967-02S1 and R01 GM130746-02S1) to E.A.

## Abbreviations

RPA: replication protein A
ssDNA: single-stranded DNA
DBD: DNA binding domain
PID: protein interaction domain
Wh: winged helix
HDX-MS: hydrogen deuterium exchange mass spectrometry

