## Supplemental Information for "Rtt105 configurationally staples RPA and blocks facilitated exchange and interactions with RPA-interacting proteins"

### Configurational stapling of RPA by Rtt105 prevents untimely recruitment of RPA-interacting proteins

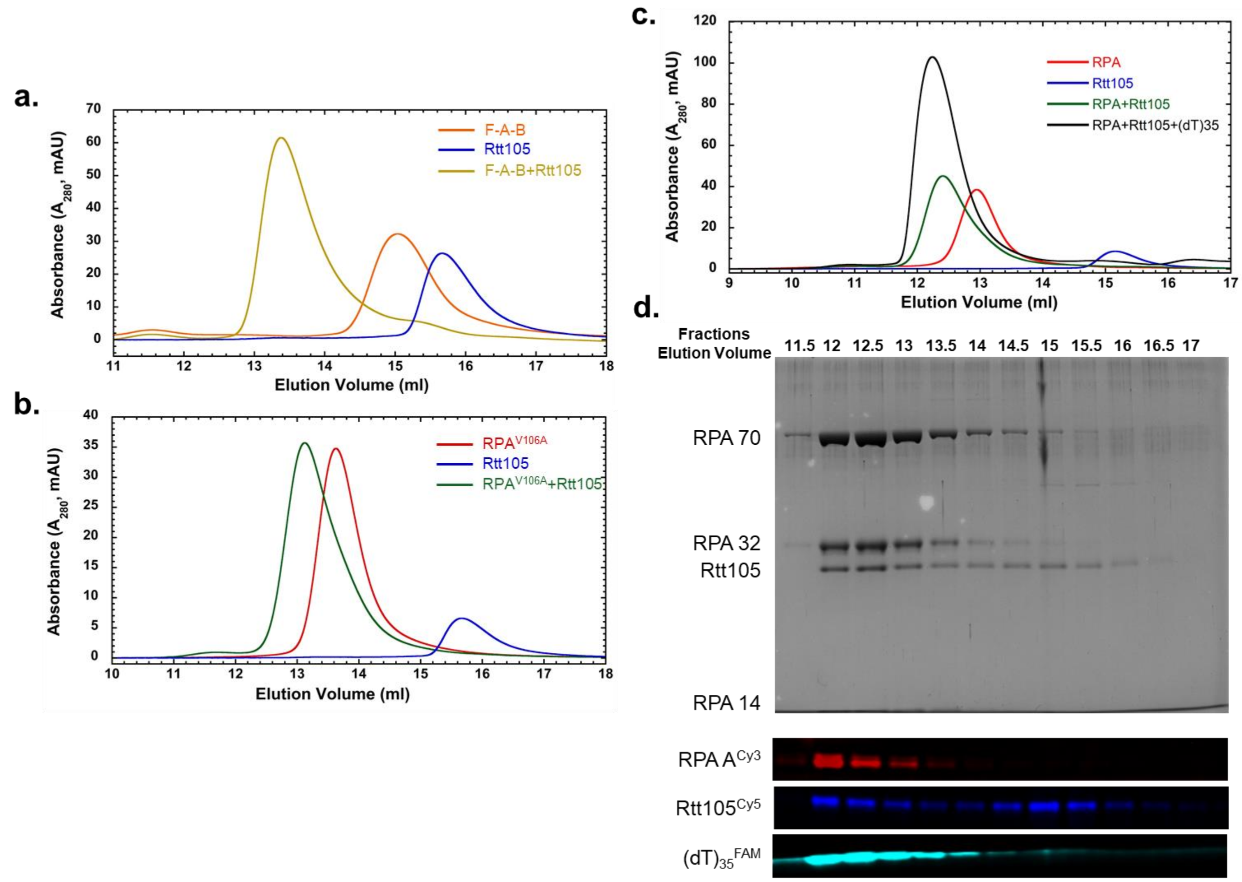


**Supplemental Figure S1: Size exclusion chromatography (SEC) analysis of complex formation between RPA and Rtt105. a)** SEC analysis of the F-A-B version of RPA (containing the OB-F, DBD-A and DBD-B domains) shows interaction with Rtt105 as observed by formation of the higher molecular weight species. **b)** A Val106 to Ala substitution in the OB-F domain of RPA70 ($RPA^{V106A}$) was predicted to not interact with Rtt105, but we observe complex formation in SEC suggesting interaction between $RPA^{V106A}$ and Rtt105. **c)** SEC analysis of 5 μM each of RPA-DBD-$A^{Cy3}$, $Rtt105^{Cy5}$, or a complex of $RPA^{Cy3}$ (5 μM) and $Rtt105^{Cy5}$ (5 μM) in presence of 5 μM 5′-FAM-$(dT)_{35}$ ssDNA. Rtt105, RPA, and the Rtt105-RPA complex migrate as single species in SEC. When ssDNA is added to this complex, a larger Rtt105-RPA-$(dT)_{35}$ complex is observed suggesting that Rtt105 remains in complex with RPA in the presence of short ssDNA oligonucleotides. **d)** Fractions from the SEC analysis in panel c were analyzed by SDS-PAGE and imaged by Coomassie staining (top) or fluorescence (bottom) using the respective excitation channels. Rtt105 migrates as a co-complex with RPA and the $(dT)_{35}$ ssDNA substrate. A small fraction of free Rtt105 (vol. 15.5 ml onwards) and the RPA-$(dT)_{35}$ complex (vol. 11.5 ml) are also observed. Representative images from three independent experiments are shown.

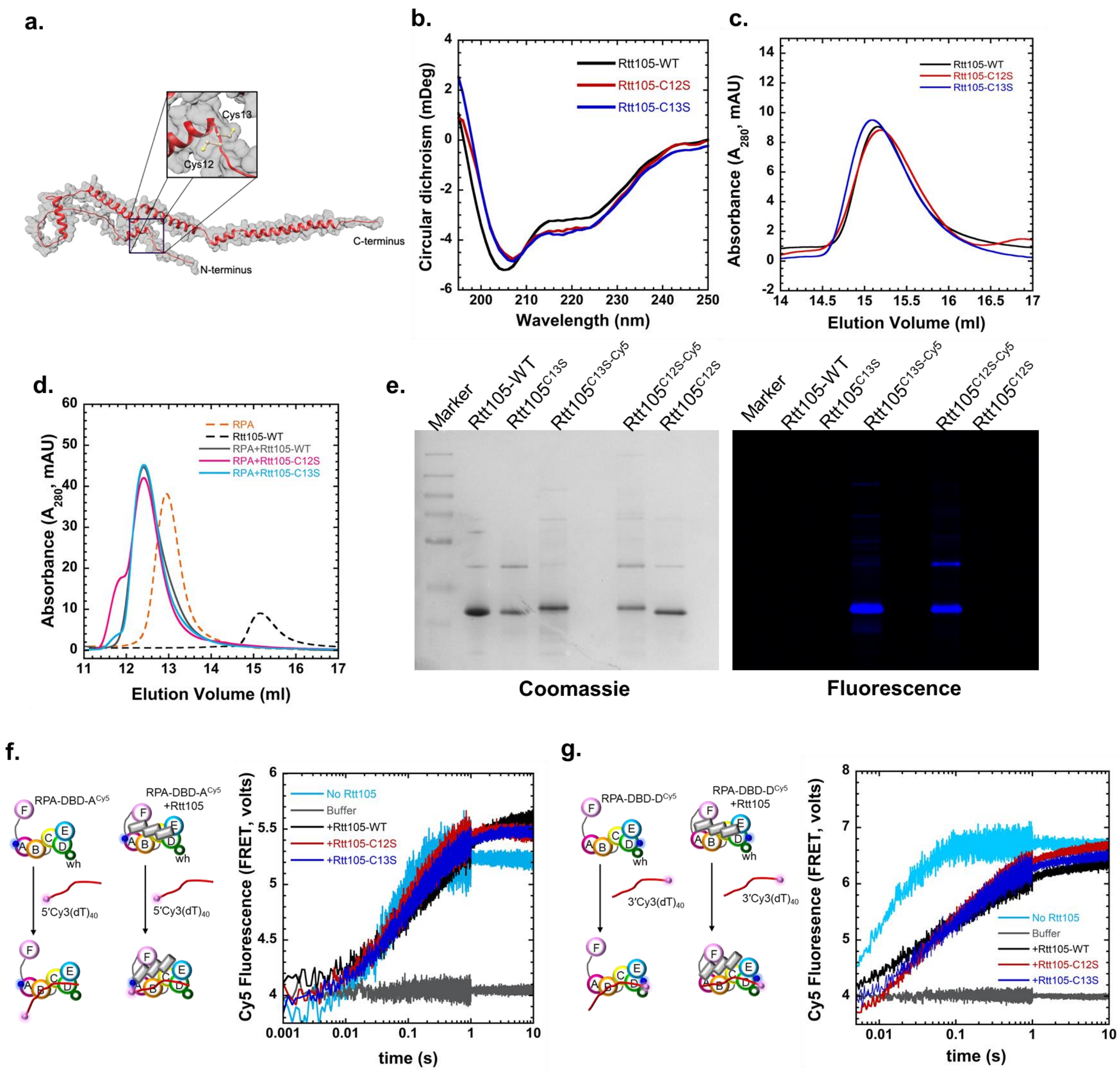


**Supplemental Figure S2. Fluorescent Rtt105 interacts with RPA similar to unlabeled Rtt105.** **a)** Positions of Cys12 and Cys13 residues are depicted in the Rtt105 structure generated using AlphaFold. Cysteine residues are represented as ball and stick on the ribbon structure (red) with a transparent surface (grey). **b)** Circular dichroism analysis of the secondary structure of Rtt105 (black) and the two single Cys variants carrying either C12S (red) or C13S (blue) substitutions show no change in the alpha helical secondary structure of Rtt105 or the mutant variants of Rtt105. **c)** Analytical size-exclusion chromatography analysis show similar elution profiles for Rtt105 and the single Cys Rtt105 variants. **d)** SEC analysis showing complex formation between RPA and Rtt105 or the single-Cys variants of Rtt105. Data show that all three proteins form stable complexes with RPA. **e)** SDS-PAGE analysis of the Rtt105, Rtt105$^{C13S}$, Rtt105$^{C13S\text{-}Cy5}$, Rtt105$^{C12S\text{-}Cy5,}$ and Rtt105$^{C12S}$ protein samples by Coomassie (left) and fluorescence (on right) imaging. **f)**

Rapid kinetic analysis of the effect of Rtt105 on RPA-DBD-A interactions with ssDNA show both Rtt105 and the single Cys variants similarly influencing the binding properties. For all Rtt105 proteins tested, the rate of binding is not affected, but the amplitudes are reduced. Cy3 placed on the $(dT)_{40}$ is excited (5′-Cy3-$(dT)_{40}$) and emission of Cy5 positioned on DBD-A (RPA-DBD-$A^{Cy5}$) via FRET were measured. Since DBD-A resides close to the 5′ end of the DNA, a high FRET signal is observed. **g)** Similar rapid kinetic experiments were used to quantitate the influence of the single Cys variants on FRET induced kinetics of Cy5 change between a 3′-Cy3-$(dT)_{40}$ substrate and RPA-DBD-$D^{Cy5}$. In this experiment, DBD-D is situated close to the 3′ end of the DNA and generates a high FRET signal. Again, no differences are observed between Rtt105 and the single-Cys variants. The rates of DNA binding are reduced, but the amplitudes are similar. Thus, substitution of either Cys residue in Rtt105 does not affect RPA binding or RPA-DNA interactions. Cys at position 12 was labeled with Cy5 to generated fluorescently labeled Rtt105. Representative images from three independent CD, SEC, and stopped flow experiments are shown.

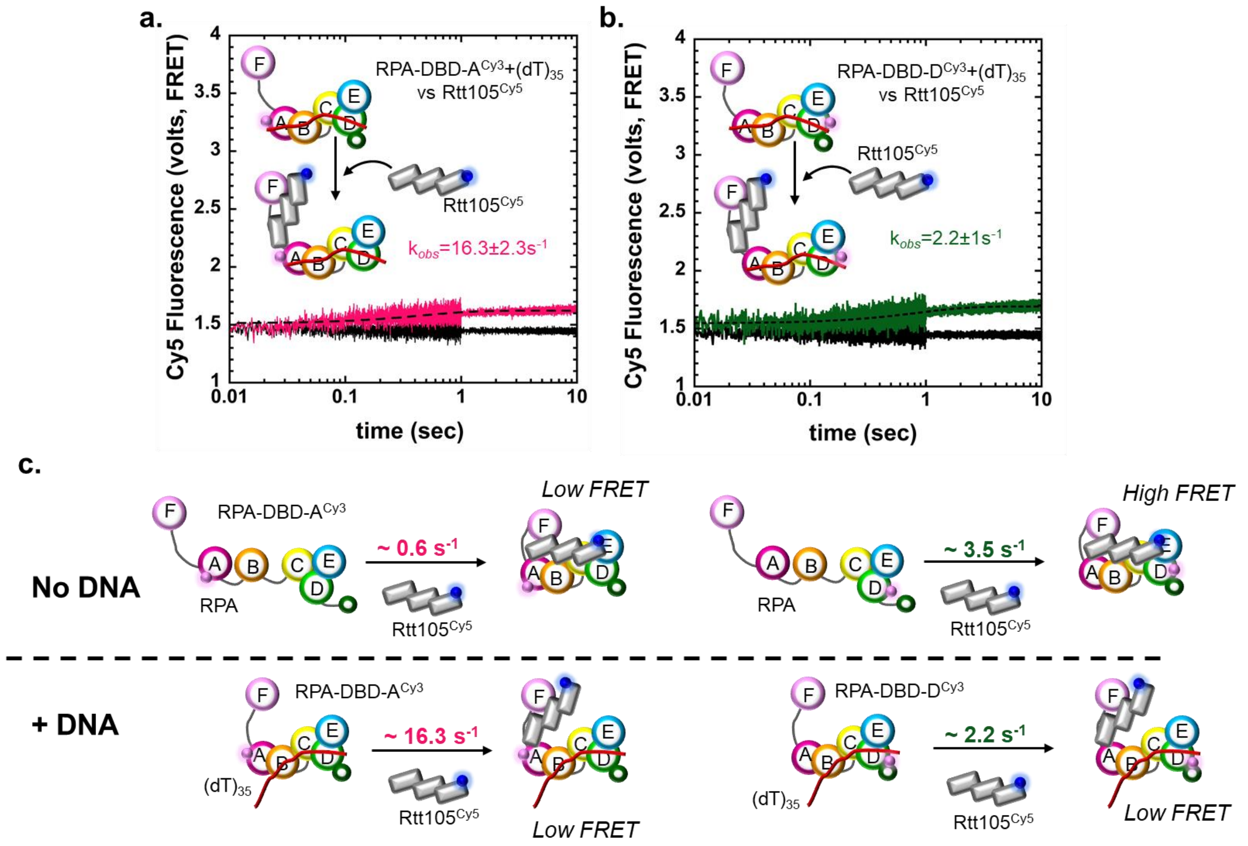


**Supplemental Figure S3: Rtt105 binds poorly to a preformed RPA-ssDNA complex.** Binding of fluorescent Rtt105 (100 nM Rtt105$^{Cy5}$) to ssDNA-bound fluorescent RPA (100 nM $(dT)_{35}$ with either **a)**100 nM RPA-DBD-A$^{Cy3}$ or **b)** 100 nM RPA-DBD-D$^{Cy3}$) was measure using stopped flow analysis. FRET between Rtt105 and respectively labeled DBDs were monitored by exciting Cy5 and monitoring the change in Cy5 fluorescence. In both cases, only a very small change in fluorescence is observed suggesting that Rtt105 binding to a preformed RPA-ssDNA complex is poor. **c)** Fitting the signals to a single step binding model revels that Rtt105 binding to DBD-D in the absence or presence of ssDNA is similar (3.5 $^{s-1}$ versus 2.5 $s^{-1}$). In contrast, the binding induced FRET signal change in DBD-A is ~25-fold faster. This suggests that Rtt105 engages closer to DBD-A faster when RPA is prebound to ssDNA. Since the F-A-B region of RPA is more dynamic and predominantly more accessible when bound to DNA it likely enables better access to OB-F and thus the increase in binding kinetics. In both panels a and b, no changes in the FRET induced Cy5 fluorescence are observed upon addition of buffer (black). A representative image from three independent stopped flow experiments is shown and the data were fit to a single exponential equation. SE from n=3 are noted.

| RTT105 Coverage | |
|---|---|
| # Amino Acids | 208 |
| Unique Peptides | 15 |
| % Coverage | 67 % |

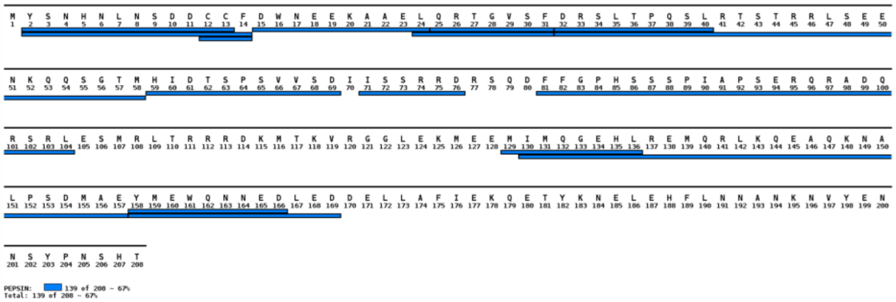


| RPA70 Coverage | |
|---|---|
| # Amino Acids | 621 |
| Unique Peptides | 77 |
| % Coverage | 65 % |

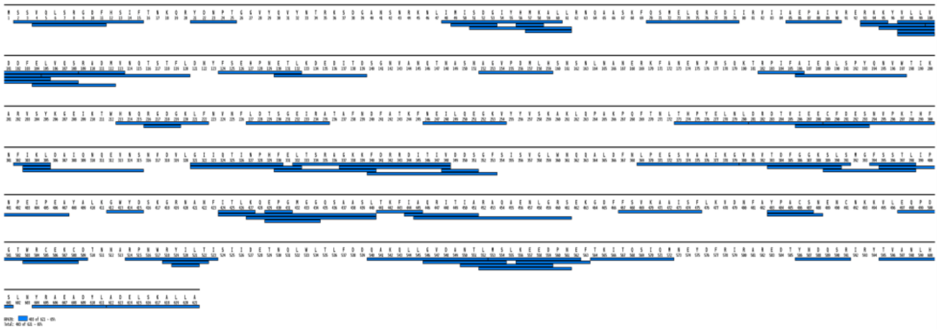

| RPA32 Coverage | |
|---|---|
| # Amino Acids | 279 |
| Unique Peptides | 16 |
| % Coverage | 30 % |

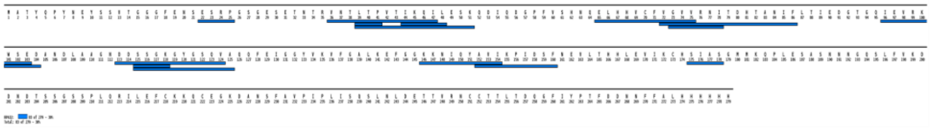

| RPA14 Coverage | |
|---|---|
| # Amino Acids | 122 |
| Unique Peptides | 8 |

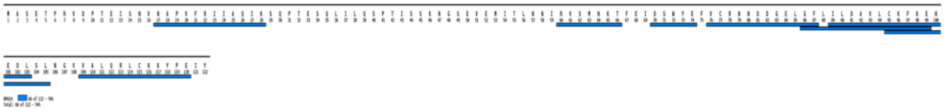

**Supplemental Figure S4. Peptide coverage in mass spectrometry analysis.** Excellent sequence coverage is observed for all three subunits of RPA and Rtt105 upon peptide digestion during mass spectrometry analysis.. The total number of amino acids, unique peptides, and percentage coverage for the individual polypeptides are denoted.

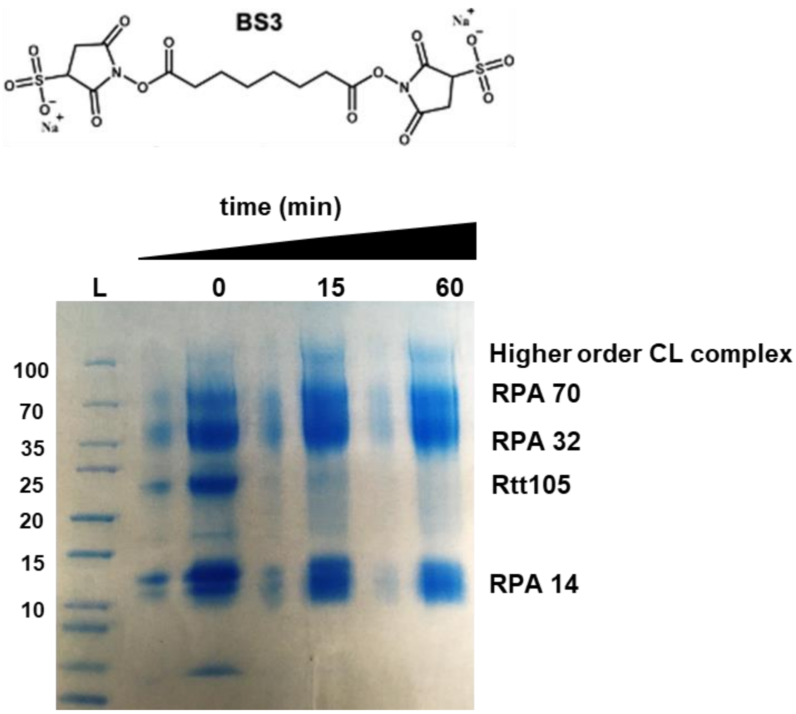


**Supplemental Figure S5. Crosslinking analysis of the RPA-Rtt105 complex.** Structure of the bis(sulphosuccinimidyl)suberate (BS3) crosslinker is shown (top). 7.75 μM RPA and 15.5 μM Rtt105 were mixed and treated with 5 mM BS3 for increasing times (0, 15 and 60 min). Samples at each time point were analyzed by SDS-PAGE and show a time-dependent increase in the crosslinking. Crosslinking of Rtt105 to RPA70 and RPA32 is readily observed with a complete shift in the Rtt105 band. The RPA14 subunit is not readily crosslinked and correspondingly, only a single crosslink between Rtt105 and RPA14 is detected in the XL-MS analysis.

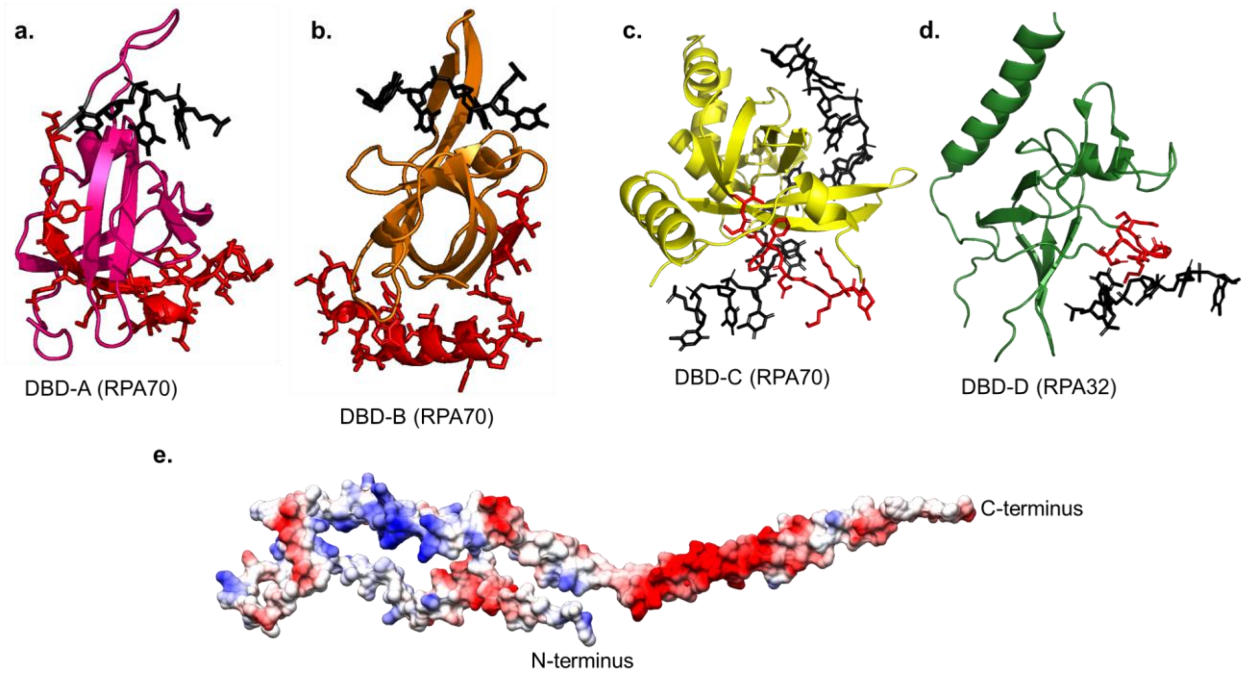


**Supplemental Figure S6. Overlap between DNA binding and Rtt105 binding regions in DBD-C and DBD-D revealed by crosslinking analysis.** The peptides identified in XL-MS are denoted as red sticks for **a)** DBD-A, **b)** DBD-B, **c)** DBD-C and **d)** DBD-D along with the bound DNA from the respective structures (black sticks). The crosslinked regions do not (or minimally) overlap with the DNA binding pockets in DBD-A or DBD-B, but significantly overlap in DBD-C and DBD-D. DBD-C and DBD-D are from the cryoEM structure of *S. cerevisiae* RPA (PDB ID: 6I52). The ssDNA-bound DBD-A and DBD-B structures were generated using SWISS-MODEL based on the DNA bound structure of *Ustilago* maydis RPA (PDB ID: 4GOP). **e)** The surface charges of Rtt105 are displayed in the AlphaFold generated model (AF-P40063-F1). Multiple pockets of negative charges are noted and can potentially bind close to the DNA binding regions in RPA.

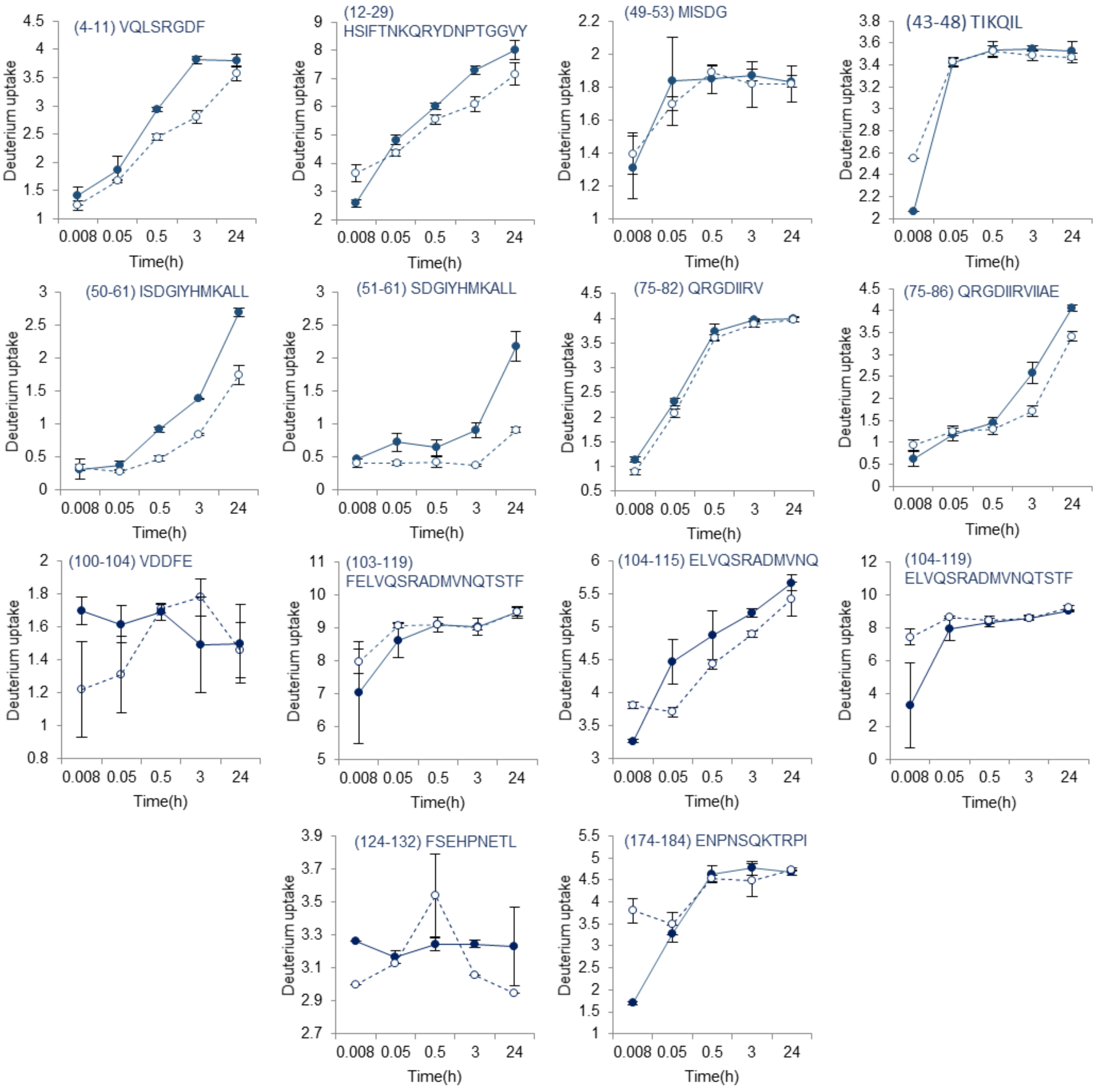


**Supplemental Figure S7. HDX-MS analysis of RPA-Rtt105 interactions and OB-F (RPA70) peptides.** HDX-MS data corresponding to specific peptides from PID$^{70N}$ (OB-F domain) are shown. Deuterium uptake for each peptide collected in the absence (●) and presence (○) of Rtt105 are plotted as a function of time. The amino acid residue number and sequence of each corresponding peptides are noted. Std. Dev. from n=3 are plotted.

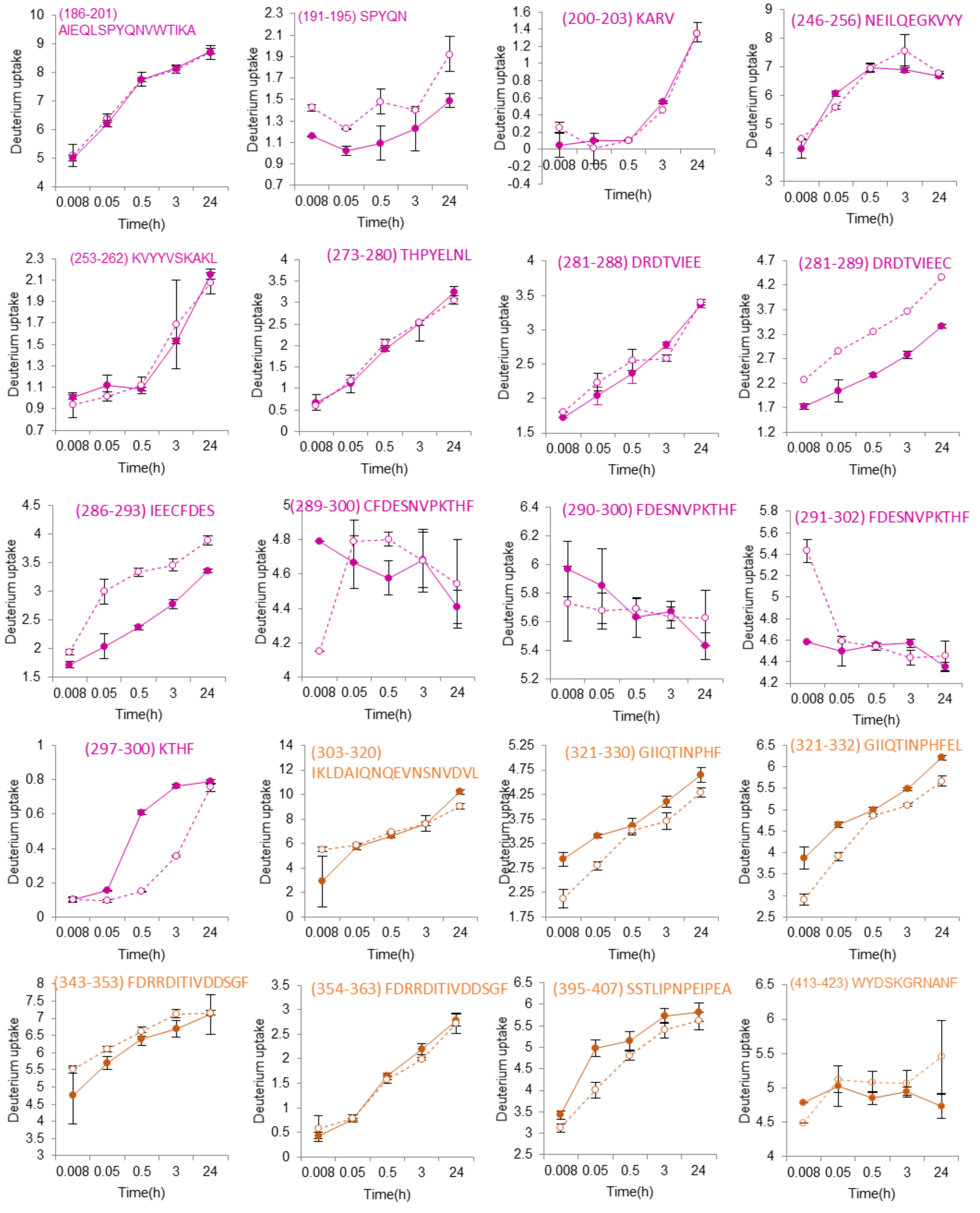


**Supplemental Figure S8**. **HDX-MS analysis of RPA-Rtt105 interactions and DBD-A & DBD-B (RPA70) peptides.** HDX-MS data corresponding to specific peptides from DBD-A (purple) and DBD-B (orange) are shown. Deuterium uptake for each peptide collected in the absence (●) and presence (○) of Rtt105 are plotted as a function of time. The amino acid residue number and sequence of each corresponding peptides are noted. Std. Dev. from n=3 are plotted.

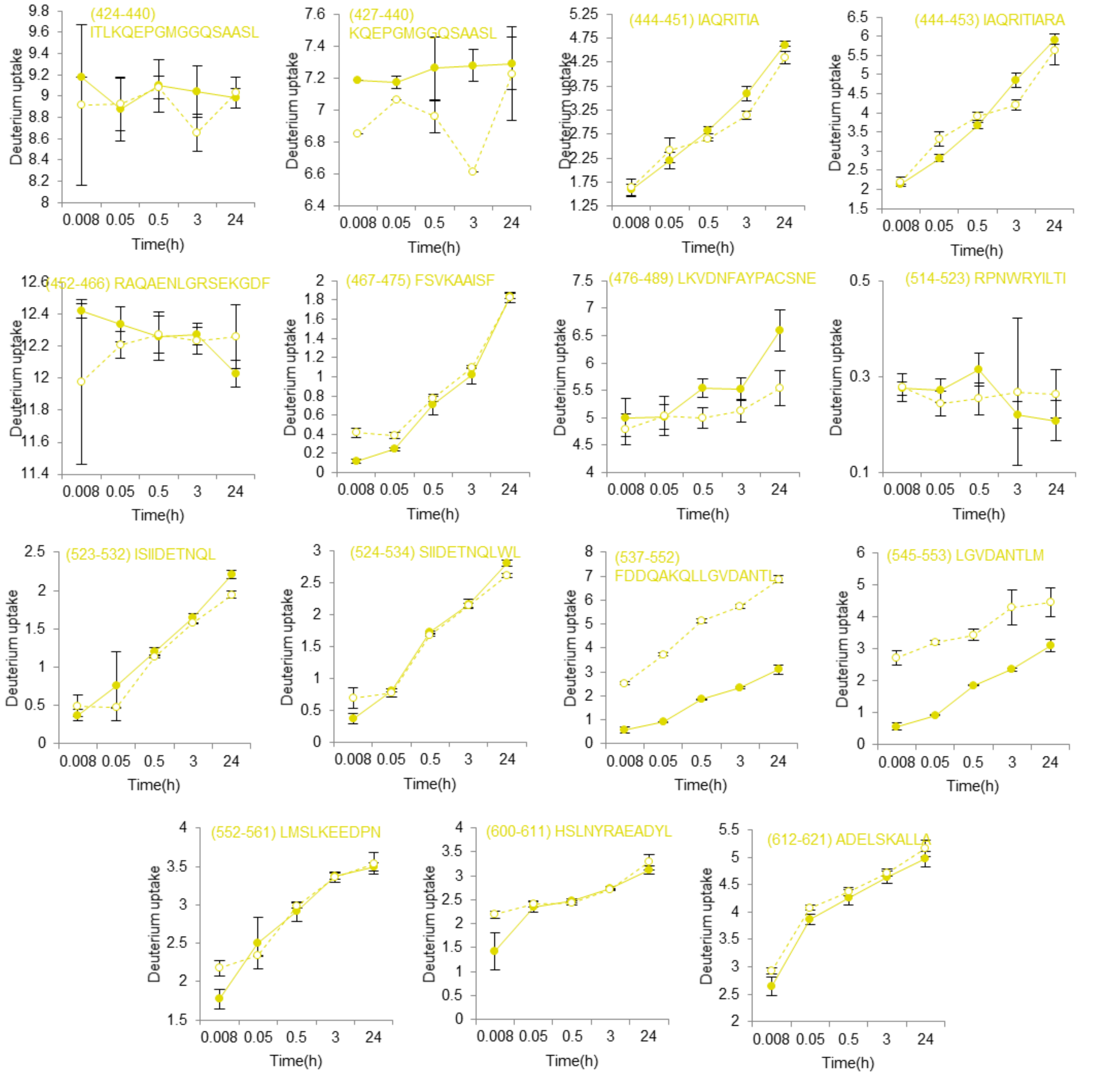


**Supplemental Figure S9**. **HDX-MS analysis of RPA-Rtt105 interactions and DBD-C (RPA70) peptides.** HDX-MS data corresponding to specific peptides from DBD-C are shown. Deuterium uptake for each peptide collected in the absence (●) and presence (○) of Rtt105 are plotted as a function of time. The amino acid residue number and sequence of each corresponding peptides are noted. Std. Dev. from n=3 are plotted.

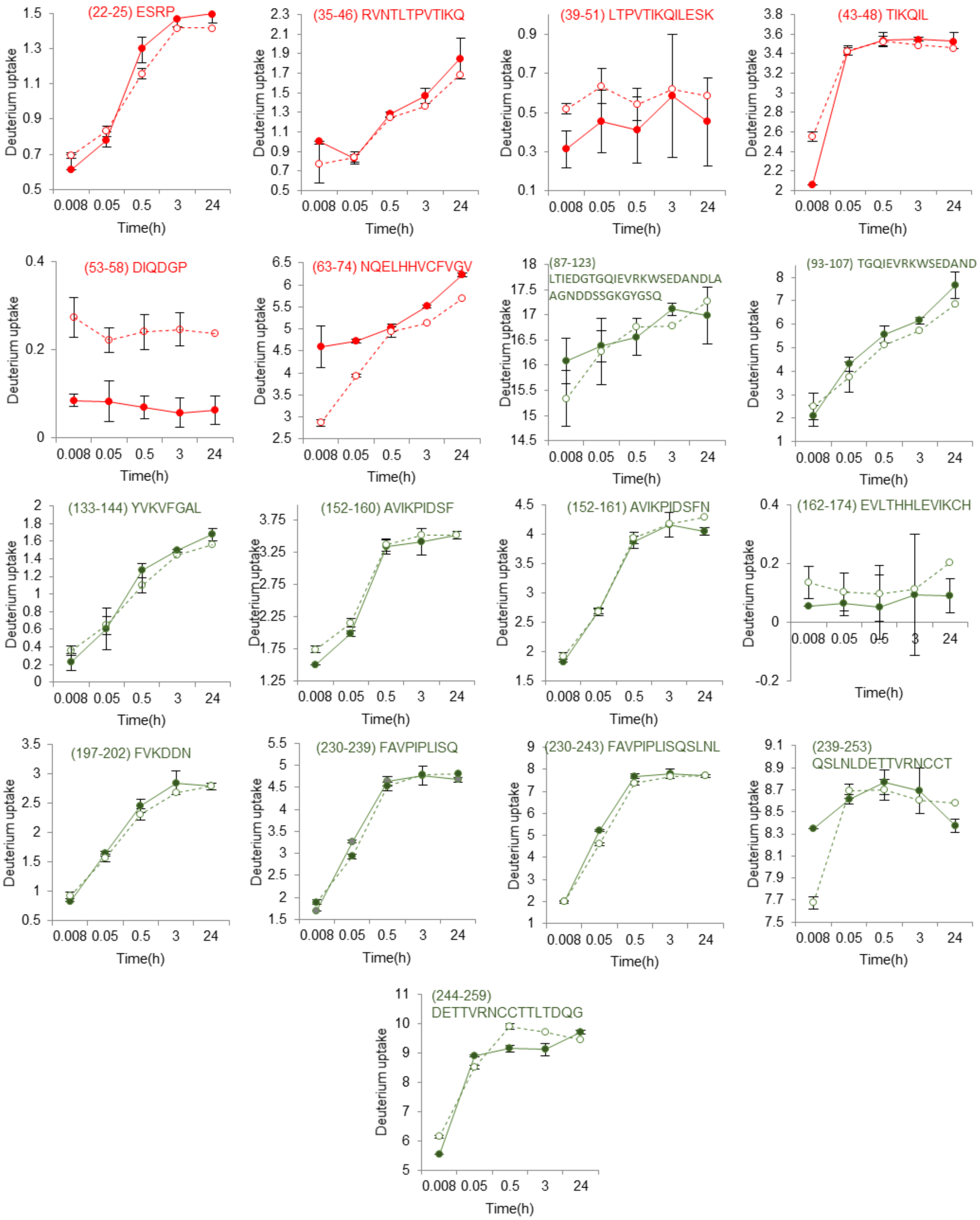


**Supplemental Figure S10**. **HDX-MS analysis of RPA-Rtt105 interactions and RPA32 (DBD-D and winged helix or PID$^{32C}$ ) peptides.** HDX-MS data corresponding to specific peptides from RPA32 are shown. The peptides in the N-terminal region of hyperphosphorylation are shown in red and DBD-D peptides are in green. Deuterium uptake for each peptide collected in the absence (●) and presence (○) of Rtt105 are plotted as a function of time. The amino acid residue number and sequence of each corresponding peptides are noted. Std. Dev. from n=3 are plotted.

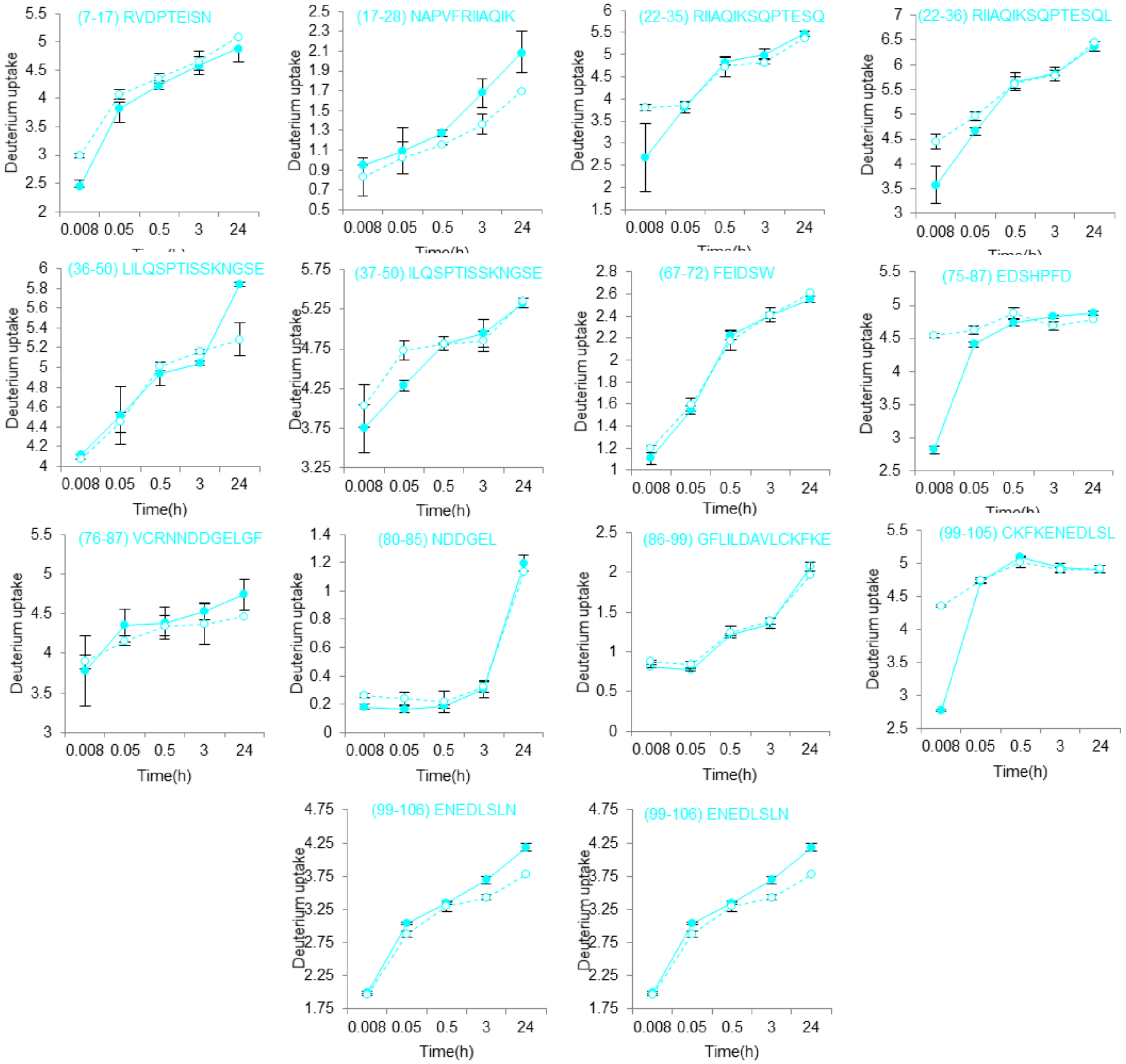


**Supplemental Figure S11**. **HDX-MS analysis of RPA-Rtt105 interactions and RPA14 (OB-E) peptides.** HDX-MS data corresponding to specific peptides from RPA14 are shown. Deuterium uptake for each peptide collected in the absence (●) and presence (○) of Rtt105 are plotted as a function of time. The amino acid residue number and sequence of each corresponding peptides are noted. Std. Dev. from n=3 are plotted.

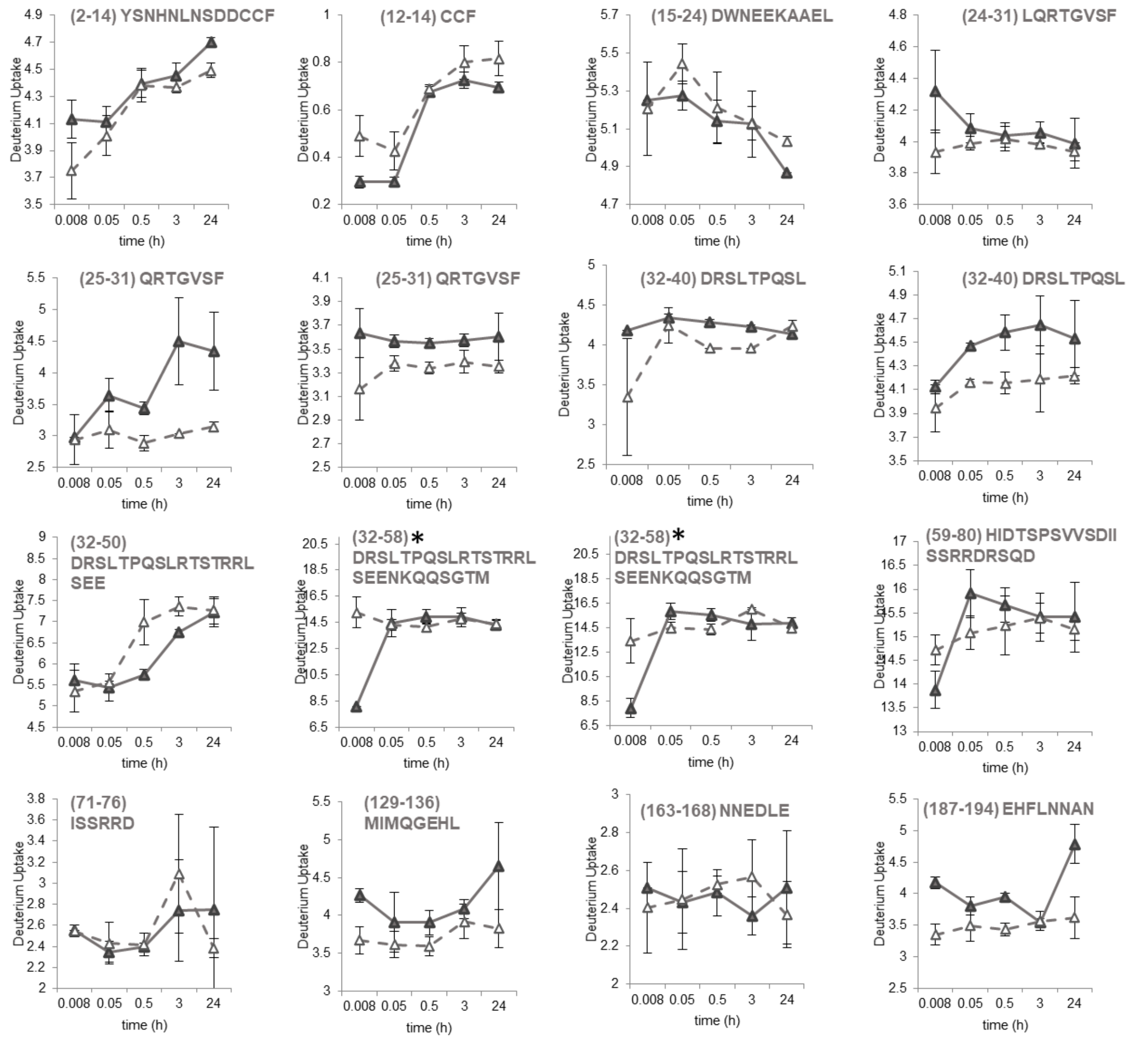


**Supplemental Figure S12**. **HDX-MS analysis of Rtt105 peptides from the Rtt105-RPA-ssDNA complex reveals DNA-induced remodeling of Rtt105.** HDX-MS data corresponding to specific peptides from Rtt105 are shown for samples measured in the absence ((▲) Rtt105-RPA) or presence of DNA ((Δ) Rtt105-RPA-$(dT)_{35}$). The amino acid residue number and sequence of each corresponding peptides are noted. Std. Dev. from n=3 are plotted. *Denotes similar peptides carrying different charge.

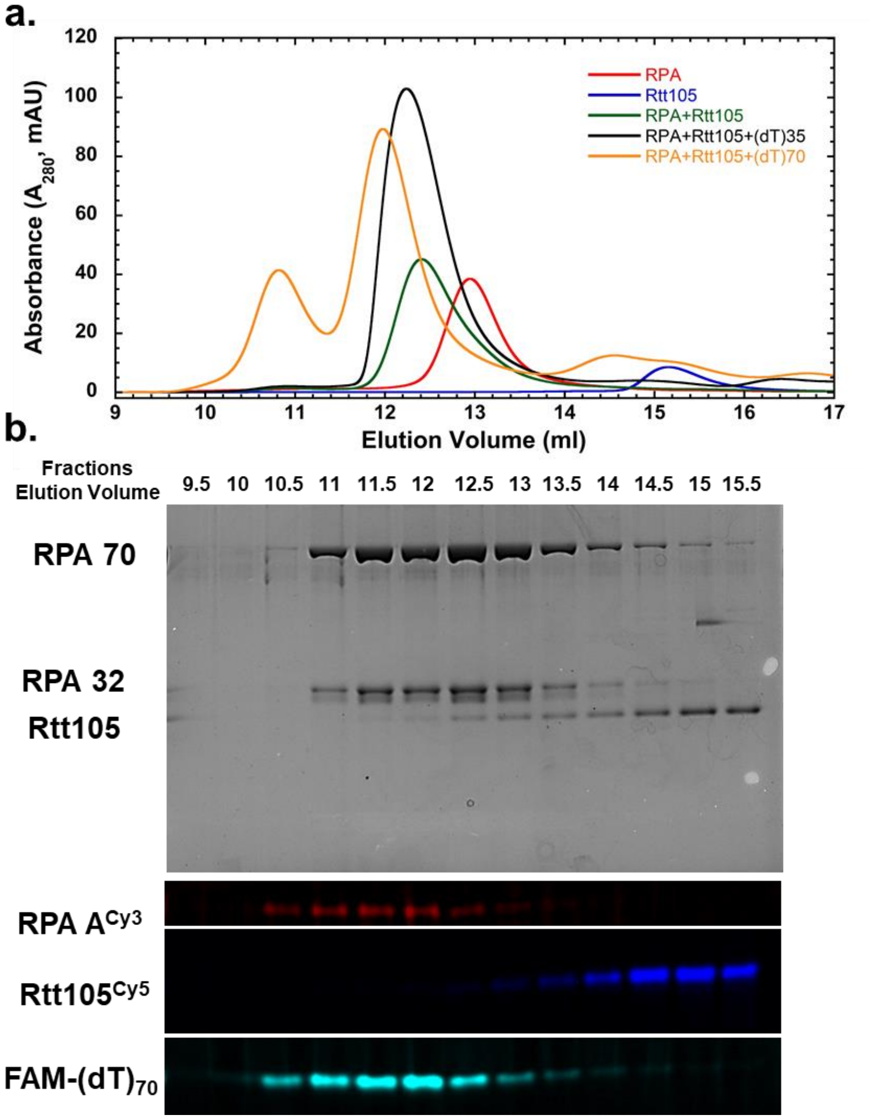


**Supplemental Figure S13. SEC analysis of Rtt105-RPA-(dT)$_{70}$ complex shows dissociation of Rtt105. a)** SEC analysis of 5 μM each of RPA-DBD-A$^{Cy3}$, Rtt105$^{Cy5}$, or preformed complex RPA$^{Cy3}$ (5 μM) +Rtt105$^{Cy5}$ (5 μM) in presence of 5 μM 5′-FAM-(dT)$_{35}$ or 5 μM 5′-FAM-(dT)$_{70}$ ssDNA oligonucleotides. **b)** The fractions from the Rtt105-RPA-(dT)$_{70}$ run in panel a were analyzed by SDS-PAGE and imaged by Coomassie staining (top) or fluorescence (bottom) using the respective excitation channels. Rtt105 dissociates from RPA in the presence of (dT)$_{70}$ ssDNA substrate. Representative images from three independent experiments are shown.

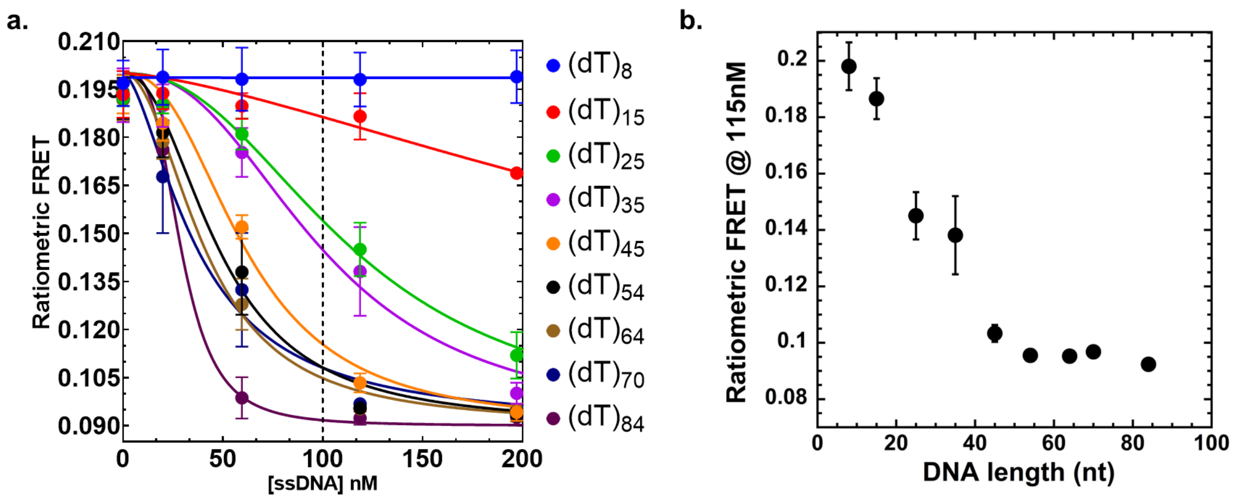


**Supplemental Figure S14: ssDNA substrates ≥35 nt in length trigger the release of Rtt105 from RPA. a)** Preformed RPA-DBD-$D^{Cy3}$-$Rtt105^{Cy5}$ complexes were mixed with increasing concentrations of $(dT)_n$ (where n = length of ssDNA) and the change in FRET was measured. **b)** Respective FRET values observed at 115 nM ssDNA concentration (~2:1 ratio of RPA:$(dT)_n$]were plotted as a function of $(dT)_n$. Increasing the length beyond $(dT)_{35}$ leads to a very low-FRET state which corresponds to Rtt105 remodeling or dissociation. The moderate FRET state (~0.14 FRET) observed around n=35 corresponds to a complex where Rtt105 is remodeled, but still bound to the RPA-DNA complex.

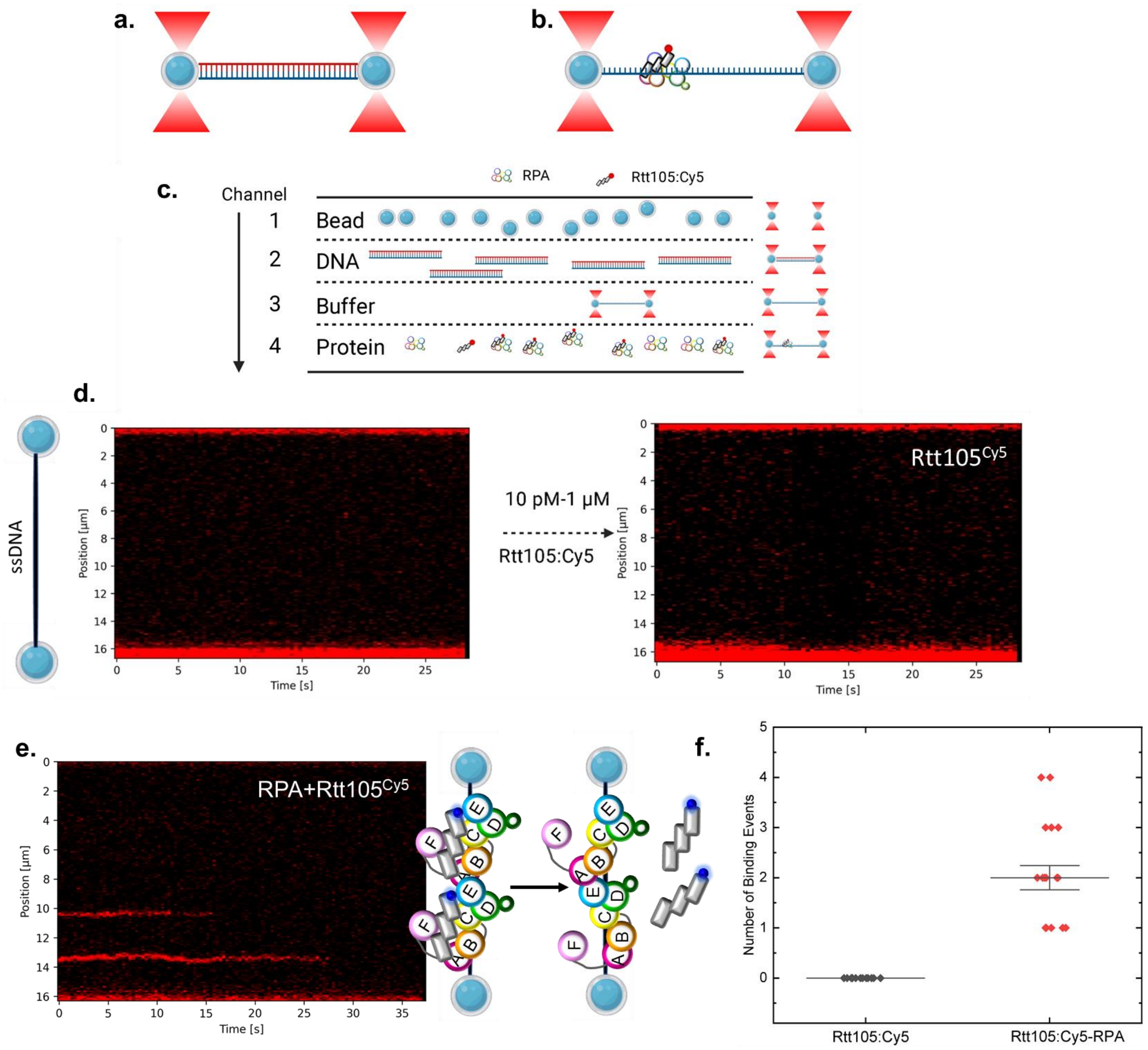


**Supplemental Figure S15. C-trap analysis shows Rtt105 poorly bound to RPA on long ssDNA**. **a)** Schematic of the C-trap setup to investigate RPA-Rtt105 interactions. Lambda dsDNA is captured between the beads and **b)** stretched to generate ssDNA. **c)** The ssDNA is the introduced into channels containing the appropriate proteins. **d)** Analysis of Rtt105$^{Cy5}$-RPA interactions on ~48.5 knt long DNA substrates reveal no Rtt105 binding to ssDNA in the absence of RPA. **e)** When the fluorescence is monitored upon exposure of the ssDNA to the Rtt105$^{Cy5}$-RPA, very few Rtt105$^{Cy5}$ binding events are observed, and these molecules also dissociate. **f)** Quantitation of the number of DNA binding events in C-trap analysis shows no Rtt105 binding to ssDNA in the absence of RPA. Roughly 2 transient binding events per RPA-coated 48.5 knt long ssDNA are observed under these experimental conditions. Data show a $t_{1/2}$ of ~5 s for the dissociation of Rtt105$^{Cy5}$ from RPA on these DNA molecules.

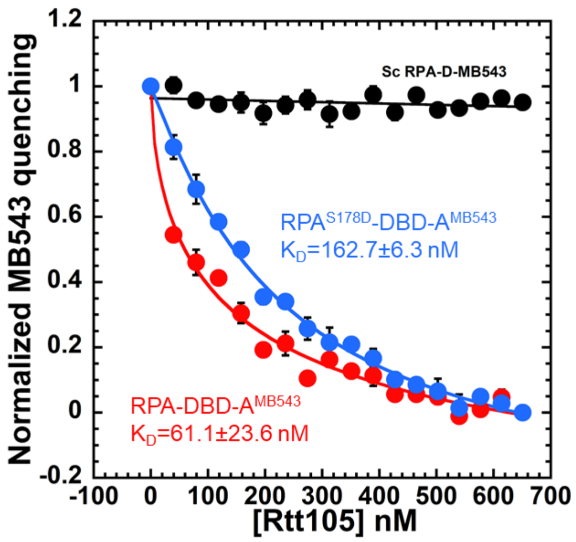


**Supplemental Figure S16. Phosphorylation of RPA at S178 reduces Rtt105 binding affinity.** Rtt105 binding to RPA or a phosphomimetic RPA$^{S178D}$ (in RPA70) was measured by following the change in fluorescence of a MB543 fluorophore positioned in DBD-A. The fluorophore, when positioned in DBD-D, does not produce a chance in fluorescence upon Rtt105 binding (black). RPA$^{S178D}$ binds with reduced affinity to Rtt105 ($K_D$=61.1±23.6 nM versus 162.7±6.3 nM for RPA and RPA$^{S178D}$, respectively.

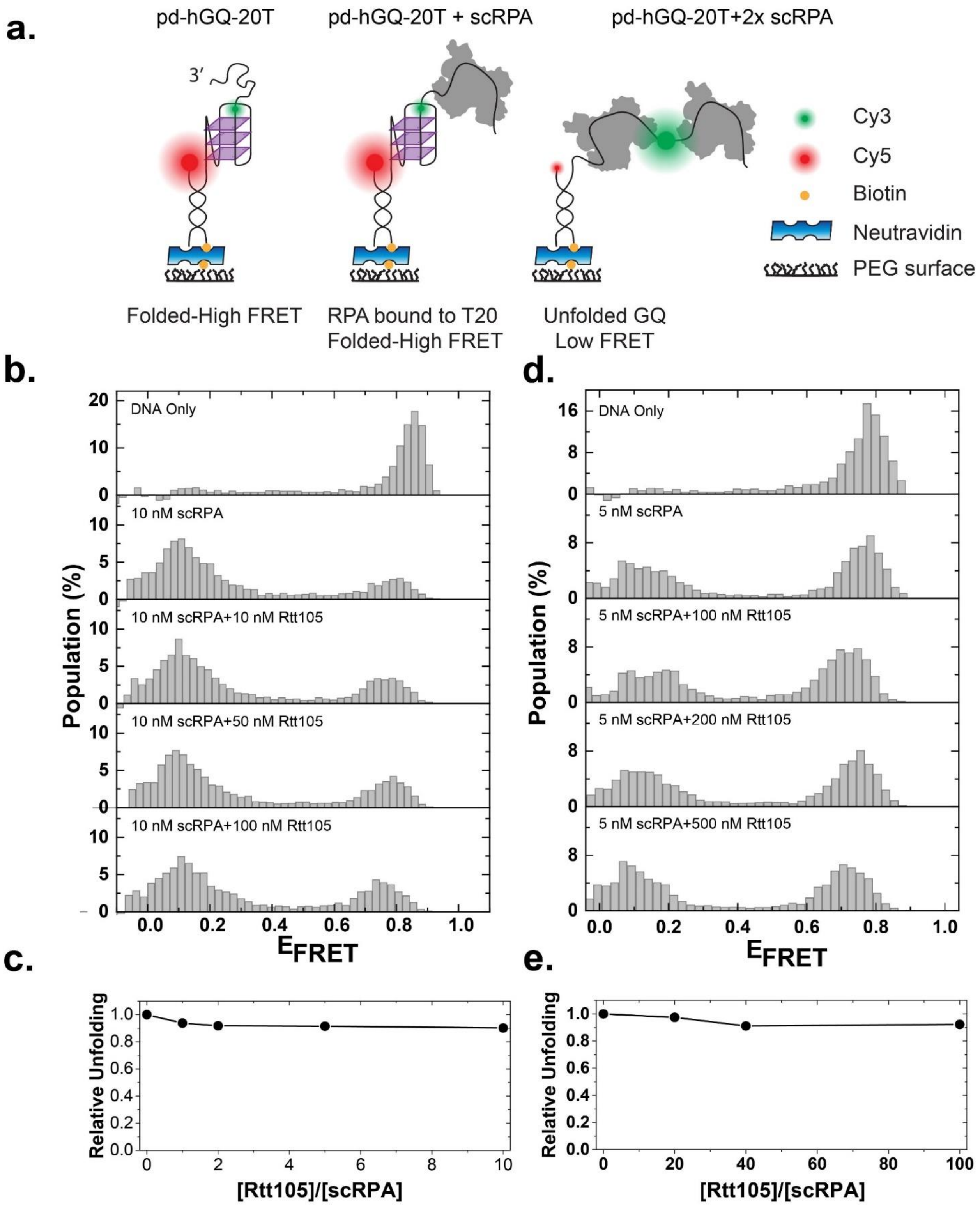


**Supplemental Figure S17. Rtt105 does not influence the G-quadruplex unwinding activity of RPA. a)** Schematics of the smFRET assay. A partial duplex DNA construct with a hGQ forming sequence and 20T overhang is immobilized on the surface of a microfluidic channel. The donor-acceptor fluorophores are placed across the hGQ. scRPA binding to the 20T overhang does not result in a FRET change until the hGQ is unfolded. Unfolding of the hGQ exposes an additional 21 nt ssDNA, which is long enough to accommodate binding of a second scRPA. Binding of scRPA to this newly exposed ssDNA results in emergence of a new low FRET peak. **b)** Top panel shows the smFRET histogram in the absence of any proteins (DNA Only) where a single high FRET peak is observed. Adding 10 nM scRPA results in ~70% of hGQ molecules being unfolded. Adding mixtures of scRPA and increasing concentrations of Rtt105 does not result in a significant change in the unfolded population. **c)** Quantification of the unfolded population for different [Rtt105]/[scRPA] ratios. The unfolded population was calculated from the cumulative population of $E_{FRET}$<0.5 states and normalized such that the 10 nM scRPA (without Rtt105) has a relative unfolding of 1.0. Before adding a new scRPA and Rtt105 mixture, the channel was washed with a high salt buffer (1 M KCl) to remove all bound proteins. The reset of the system was confirmed by restoring the histogram with a single high FRET peak after adding imaging buffer at original salt concentration (50 mM KCl+2 mM $MgCl_2$). **d)** and **e)** Similar measurements and analysis as those performed for (b) and (c) when scRPA concentration was kept at 5 nM and higher [Rtt105]/[scRPA] ratios were reached. Even at [Rtt105]/[scRPA]=100, the unfolded population was within 10% of its original value.

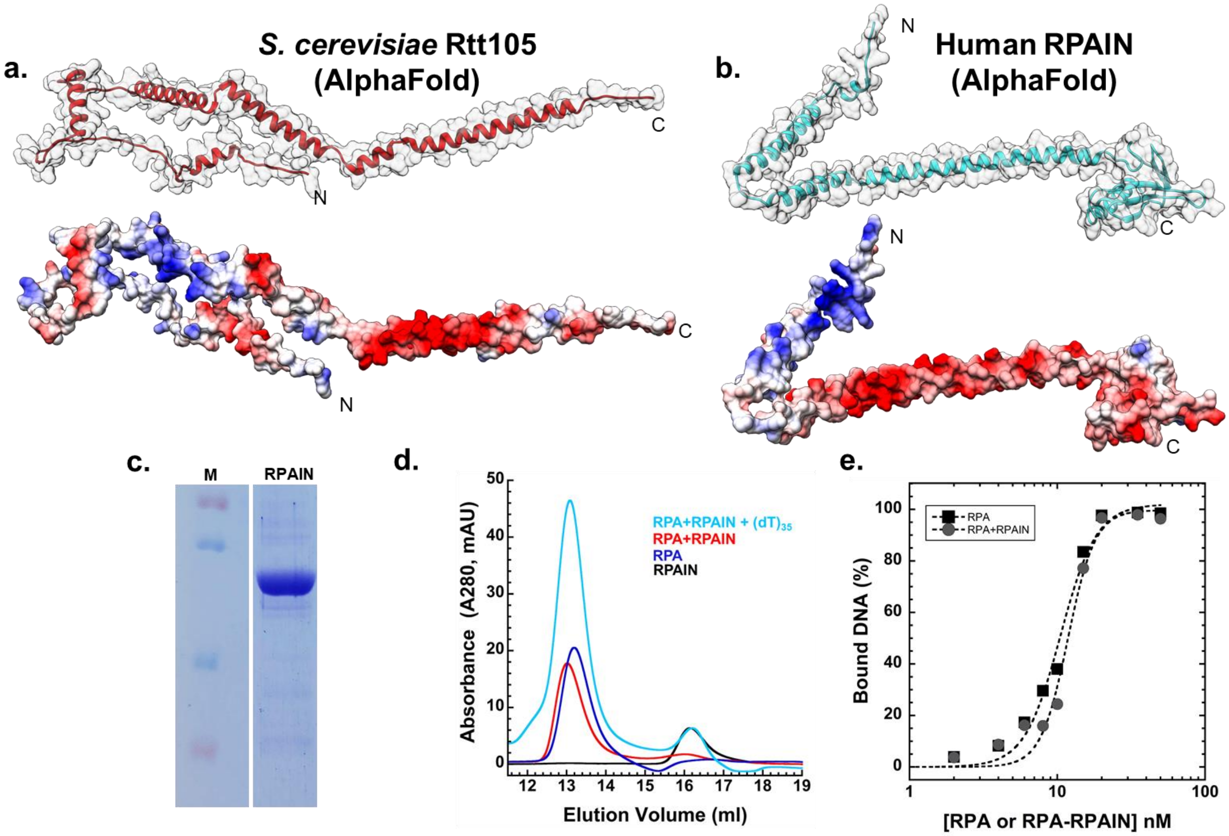


**Supplemental Figure S18. Rtt105 does not enhance the ssDNA binding activity of RPAIN, the human ortholog of Rtt105.** AlphaFold generated models of **a)** ScRPA (AF-P40063-F1) and **b)** human RPAIN (AF-Q86UA6-F1). The electrostatic maps of both structures are displayed as well and show regions of strong negative charge. **c)** SDS-PAGE analysis of recombinantly purified RPAIN. **d)** SEC analysis of 3 μM hRPA (blue) and 3 μM RPAIN (black) show singly migrating species of each protein. The RPA-RPAIN complex generated by mixing 3 μM of each protein (red) migrates as a larger complex and shows interaction between the two proteins. Addition of 3 μM $(dT)_{35}$ to the hRPA-RPAIN complex results in the release of RPAIN from RPA (cyan). **e)** The ssDNA binding activity of hRPA was measured by analyzing complex formation in EMSA analysis in the absence or presence of RPAIN. Experiments were performed using 20 nM 5′-Cy5-$(dT)_{30}$ ssDNA and titrating increasing concentrations of hRPA or the hRPA-RPAIN complex. Plot of the percent DNA bound as a function of protein concentration shows no RPAIN influence of the DNA binding activity of hRPA ($K_D$ = 10.52 nM and 11.8 nM in the absence or presence of Rtt105, respectively).
